# Hypoxia-Induced Metabolic Reprogramming and Markings of Cell Fate in Concentric Arterial Hypertrophy

**DOI:** 10.1101/2025.07.09.663881

**Authors:** Lucas Ferreira de Almeida, Jason P. Smith, Manako Yamaguchi, Silvia Medrano, Alexandre Martini, Daisuke Matsuoka, Zuzanna J Juśkiewicz, Brant E Isakson, Hiroki Yamaguchi, Dilza Trevisan Silva, Thomas Wagamon, Maria Luisa S. Sequeira-Lopez, R. Ariel Gomez

## Abstract

Chronic inhibition of the renin-angiotensin system (RAS), while widely used to treat hypertension, can lead to an underrecognized form of vascular disease marked by concentric arteriolar and arterial hypertrophy (CAAH). Here, using two lineage-traced mouse models of genetic renin deletion and sustained RAS blockade, we uncover a pathogenic cascade initiated by renin-lineage cell fate reprogramming. Loss of endocrine identity and transformation of smooth muscle cells drives a shift toward a fibrotic, inflammatory, and secretory phenotype that remodels the extracellular matrix and promotes vascular thickening and luminal narrowing. Integrated transcriptomic, proteomic, and metabolomic profiling revealed a hypoxia-linked metabolic switch—characterized by succinate accumulation and NAD^+^ depletion—coupled to Hif activation and disease progression. We identify Cdh13 and collagens (including Col1a1 and Col12a1) as early urinary biomarkers and define a 10-gene molecular signature of CAAH with potential clinical application. These findings establish renin-lineage cell plasticity and metabolic dysfunction as central drivers of CAAH and nominate candidate biomarkers for early detection and therapeutic targeting in RAS-inhibited patients.

## Introduction

Over 1.3 billion people worldwide suffer from hypertension including 30% of the US population. Alarmingly, these numbers continue to rise^1,2^. The problem is not limited to adults, affecting approximately 2-5 % of children representing 3-5 million US children^3^. Because hypertension is a major risk factor for cardiovascular disease, stroke, heart and renal failure, the American Heart Association and other health organizations have developed stricter blood pressure (BP) guidelines suggesting lowering the threshold limits to treat elevated BP^4,5^. Because of their effectiveness, renin-angiotensin system (RAS) inhibitors are used frequently as the first line of treatment. However, the inhibition of the RAS causes profound changes in renin cells and the renal arterial tree of young and adult animals, including humans^6^. Experimental or spontaneous mutations of any of the renin-angiotensin system (RAS) genes or treatment with RAS inhibitors in all mammals examined, including humans, leads to concentric arterial and arteriolar hypertrophy (CAAH) of the kidney vasculature^6–9^. Through previous efforts, we established that renin cells *per se* are required for the vascular disease to develop, and that under constant stimulation renin cells become hypertrophic, recruit smooth muscle cells, and together accumulate along the walls of arteries and arterioles in a disordered pattern^7,10^. Knowledge of this silent, progressive, and severe vascular disease remains limited. We hypothesized that under constant, unrestrained stimulation (by the combination of lower blood pressure and/or lack of Angiotensin II suppressive feedback) these hypertrophic renin cells develop loci-specific -chromatin and transcriptomic-changes that transform them from myoepithelioid cells located at the juxtaglomerular (JG) tips of the afferent arterioles into embryonic-matrix-secretory cells spread throughout the renal arterial tree. Because the transformed renin cells induce the concentric accumulation of numerous and immature arteriolar SMCs leading to the narrowing of vascular lumens, we further hypothesized that this phenomenon would yield hypoxic microenvironments in the kidney. The underlying mechanism driving CAAH and potential noninvasive biomarkers for this pathology remain largely unknown. Using genetically modified mice harboring fluorescent lineage tracers, we compared the physiology, morphology, and genomic changes following deletion of the renin gene or long-term inhibition of angiotensin converting enzyme using captopril, a common medication to treat hypertension in humans. Using both models, we characterized the histopathological changes that occurred during the development and progression of CAAH. We next investigated whether renin-lineage cells underwent progressive and loci-specific changes in their transcriptional or chromatin profiles associated with CAAH. Based on our previous findings, we tested whether renin and smooth muscle cells synthesized factors that modified the composition of the extracellular milieu responsible for the arterial disease. Next, because our mouse models have 100% penetrance of the arteriolar disease, we further posited that the unique secretome released by the reprogrammed renin and immature SMC population would generate traceable urinary and plasma biomarkers that mirror the emergence and trajectory of this vascular pathology. This study reveals a novel paradigm in which chronically stimulated renin-lineage cells act as drivers of a maladaptive pathological arteriolar remodeling and offers a path toward biomarker-guided detection and monitoring of this silent but progressive vascular disease.

## Results

### Chronic RAS suppression results in concentric arterial and arteriolar hypertrophy (CAAH) and renal functional impairment

To delineate the mechanisms driving CAAH under chronic RAS suppression, we used two complementary murine models: genetic ablation of renin (*Ren1*^*c-/-*^;*Ren1*^*cCre*^;*R26R*^*mTmG*^; *Renin*^*null*^/*Ren1*^*c*^*KO* Fig. 1A) and long-term pharmacological inhibition of the RAS using captopril (*SMMHC*^*CreERT2*^;*R26R*^*tdTomato*^;*Ren1*^*cYFP*^ mice; Fig. 1B). Both models allowed longitudinal analysis of vascular remodeling and renin-lineage cell dynamics across 1, 3, and 6 months.

Deletion of the renin gene or RAS inhibition using captopril resulted in the progressive thickening of the renal arterial tree, marked by concentric hypertrophy of interlobular arteries and afferent arterioles, accompanied by luminal narrowing, perivascular fibrosis, and mononuclear cell infiltration (Fig. 1C-D). *α*-SMA staining revealed aberrant expansion of immature smooth muscle cells (SMCs) within the vascular media (Fig. 1C-D). GFP^+^ *Ren1*^*c*^*KO* cells or YFP^+^ cells from *SMMHC*^*CreERT2*^;*R26R*^*tdTomato*^;*Ren1*^*cYFP*^ mice accumulated along the vessel walls in chaotic, disorganized manners and contributed directly to mural thickening (Fig. 1C-D). Three dimensional visualization confirmed extensive remodeling, with hypertrophic vessels encased by GFP^+^/YFP^+^ cells extending far beyond the juxtaglomerular niche, replacing the physiological distribution of renin-lineage cells (Fig. 1C-D and supplementary videos,,, and). In captopril-treated mice, these changes intensified with treatment duration and were absent in untreated controls, in which renin expression remained confined to the JG area (Fig. 1D).

**Fig. 1.**
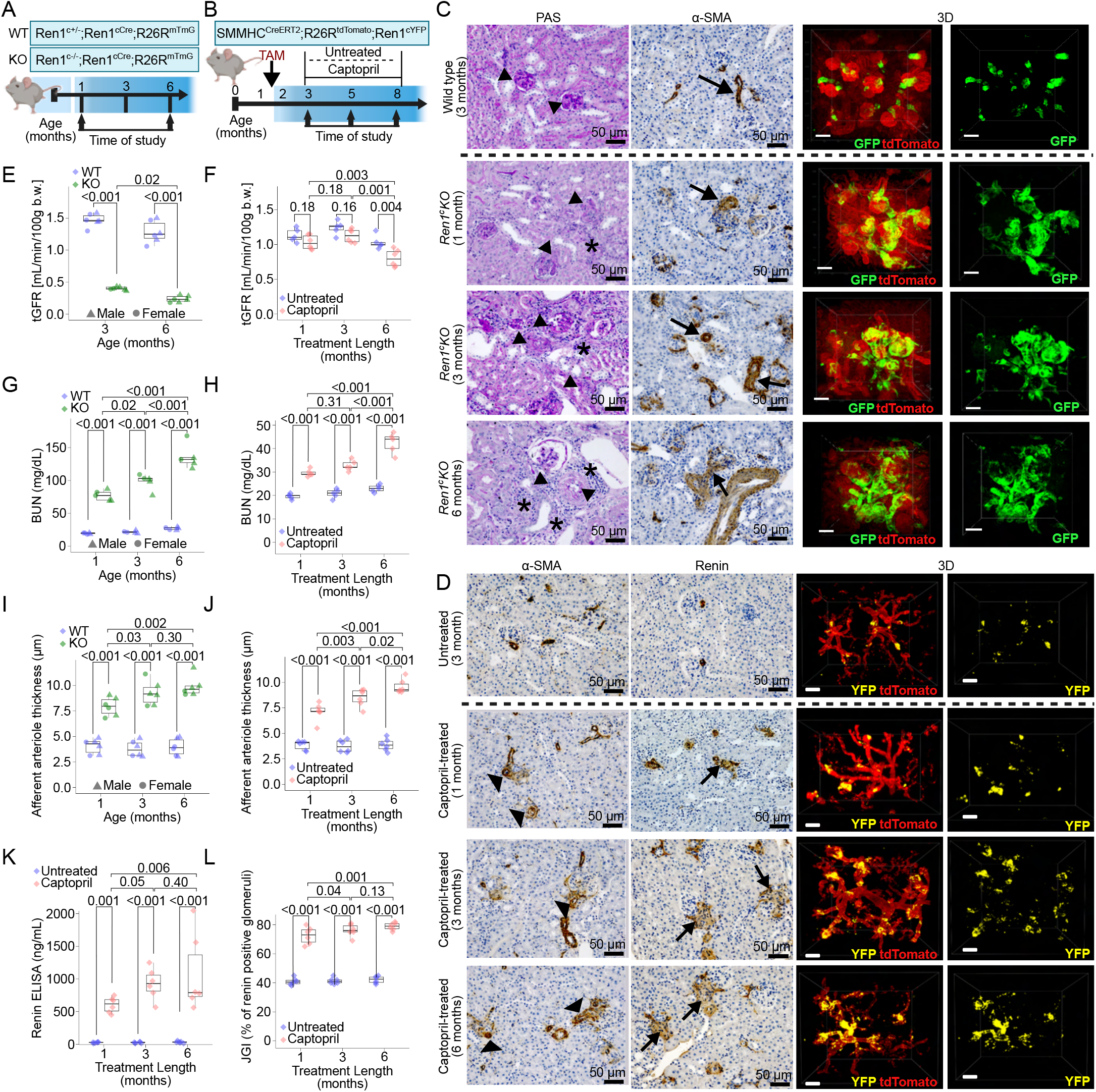
Chronic RAS suppression leads to progressive renal functional impairment. Two mouse models of CAAH were utilized: (A) genetic ablation of Ren1 (Ren1^c^KO) and (B) chronic RAS inhibition using captopril. (C) Representative images of kidney cortex showing Periodic Acid-Schiff (PAS), α-SMA staining, and 3D visualization of the Ren1 ablation mouse model demonstrated progressive vascular remodeling in the kidney (arrowheads in control group show normal architecture of periglomerular arterioles with a single layer of smooth muscle cells (SMCs); arrows in knockout animals show concentric hypertrophy and narrowed lumens; asterisks represent mononuclear cell infiltration; scale bar 50 µm). (D) α-SMA, Ren1 staining, and 3D visualization of the RAS inhibition mouse model recapitulates the findings from the genetic ablation model (arrowheads show concentric hypertrophy of SMCs; arrows indicate Ren1 abundance; scale bar 50 µm). Transdermal glomerular filtration rate (tGFR) was significantly reduced in Ren1^c^KO mice within 3 months (E) and the captopril-treated mice by 6 months (F). Blood urea nitrogen (BUN) levels were significantly elevated within 1 month in Ren1^c^KO mice (G) and in captopril-treated mice (H). The mean wall thickness of afferent arterioles was significantly thicker in Ren1^c^KO mice (I) and the captopril-treated mice (J) over time. (K) Circulating renin was significantly elevated in captopril-treated mice within 1 month of treatment. (L) The juxtaglomerular index (JGI) was significantly increased after 1 month of captopril treatment. All the experiments were performed with 6 animals per group. In the Ren1 genetic ablation model, triangle points are male and circles are female mice. The chronic RAS inhibited mouse model produces only male study animals (diamonds). All data are reported as means ± standard deviation.

Functionally, CAAH was associated with progressive decline in renal function. Glomerular filtration rate (GFR), measured by FITC-sinistrin clearance, was significantly reduced in *Ren1*^*c*^*KO* mice at 3 and 6 months, and in captopril-treated mice after 6 months (Fig. 1E-F). Blood urea nitrogen (BUN) levels were significantly elevated in both models (Fig. 1G-H), confirming the onset of kidney dysfunction despite effective systemic BP control in captopril-treated animals (Fig. S1). Further, the afferent arterioles progressively thickened over time in both animal models (Fig. 1I-J).

Together, these findings revealed that sustained RAS suppression—whether genetic or pharmacologic—induces a stereotyped cascade of vascular remodeling, initiated by renin-lineage cell reprogramming and SMC expansion. This remodeling leads to progressive vascular occlusion, inflammation, and fibrosis, ultimately compromising renal perfusion and function. The observed spatiotemporal convergence of histopathological and functional changes underscores a pathogenic mechanism with translational relevance for hypertensive patients under chronic RAS blockade.

### Evaluation of urinary protein reveals potential early biomarkers of CAAH

Given the progressive vascular remodeling, renin cell reprogramming, and renal dysfunction observed in both genetic and pharmacologic models of chronic RAS suppression (Fig. 1), we next asked whether secreted urinary proteins could serve as early, non-invasive biomarkers of CAAH onset and progression. To address this, we performed untargeted, osmolality-normalized proteomic profiling of spot urine samples from *Ren1*^*c*^*KO* and control mice (phenotypically *Ren1*^*c*^*WT*) at 1 and 6 months of age (Fig. 2A). At 1 month, 158 proteins were detected, including 32 unique to *Ren1*^*c*^*KO* mice. By 6 months, the urinary proteome expanded to 174 proteins, with 30 exclusive to *Ren1cKO* mice and 10 to controls (Fig. 2A).

**Fig. 2.**
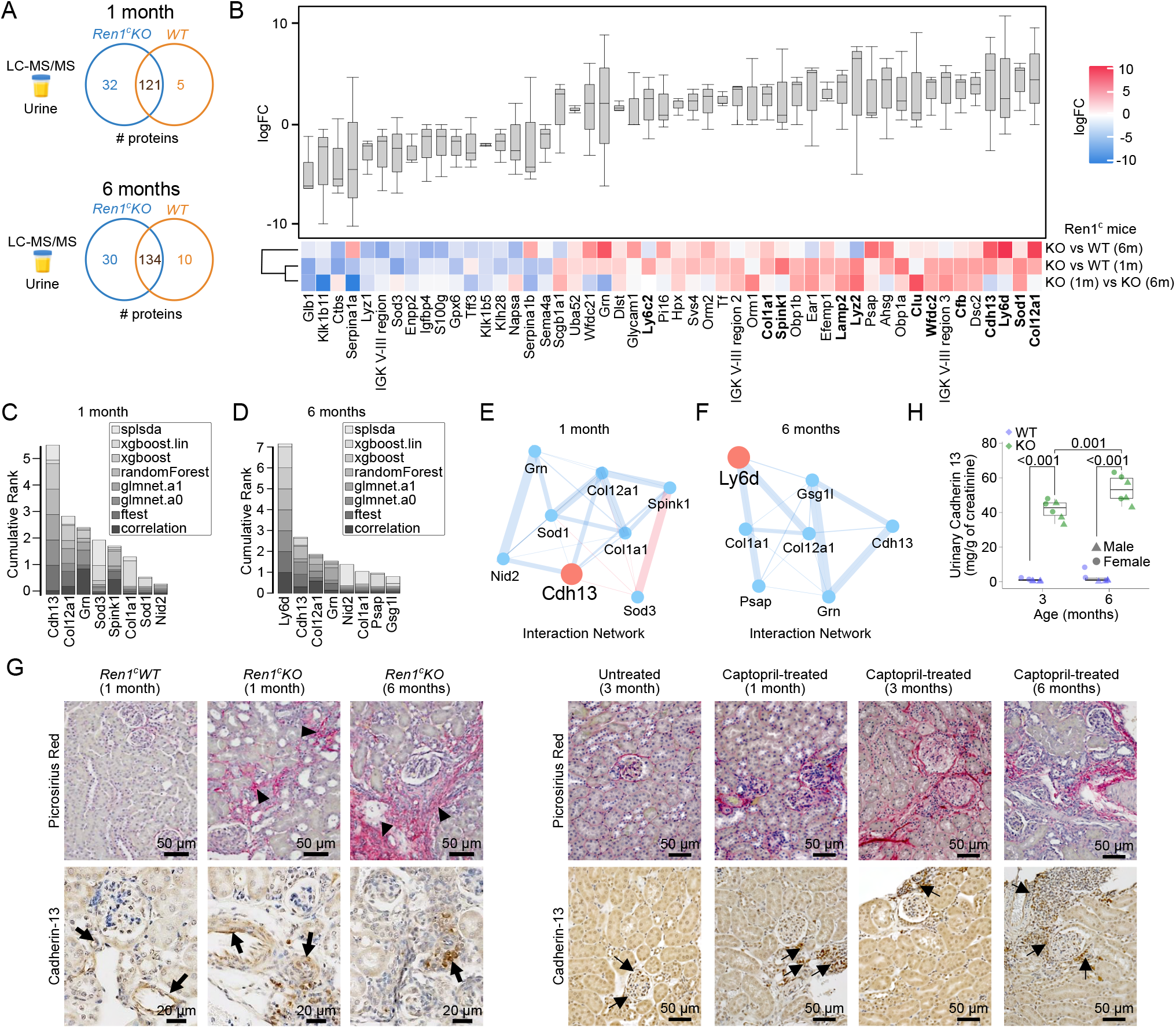
A distinct urinary protein signature in mice with CAAH was established using Ren1^c^KO mice. (A) Venn diagram showing the number of proteins detected comparing Ren1^c^KO mice vs controls (i.e. phenotypically WT) mice at 1 and 6 months of age. (B) Proteomic connectivity heatmap of KO and WT mice at 1 and 6 months, showing clustering based on profile similarity. Potential biomarkers were identified using the cumulative rank of multiple machine learning models at (C) 1 month and (D) 6 months of ages. (E) At 1 month of age, Cdh13 (cadherin 13) was the top ranked candidate and interaction network analysis revealed positive interactions with collagens. (F) At 6 months of age, Ly6d was the top candidate and network analysis identified positive interactions with both collagens and cadherin 13. (G) (upper) Picrosirius red staining revealed increased collagen deposition (arrowhead) in both Ren1^c^KO and captopril-treated kidneys, particularly in hypertrophic vessels and inflammatory infiltration regions. (lower) Cdh13 immunostaining identified elevated vascular Cdh13 abundance (arrow) in Ren1^c^KO and captopril-treated kidneys. (H) Urinary Cdh13 levels, measured by ELISA at 3 and 6 months of age, were exclusively detected in KO mice. Triangle points are male and circles are female mice. All data are reported as means ± standard deviation. n=6 animals per group.

Proteins associated with extracellular matrix remodeling, oxidative stress, and inflammation—such as Col12a1, Col1a1, Cdh13, Grn, Sod1, Sod3, Spink1, and Ly6d were significantly enriched in *Ren1*^*c*^*KO* urine (Fig. 2B). To prioritize candidate biomarkers, we applied machine learning algorithms — sparse partial least squares (Spls), elastic net, random forests, and extreme gradient boosting (XGBoost) — and generated a cumulative feature ranking (Fig. 2C-D). At 1 month, Cdh13, Col12a1, Grn, Col1a1, Spink1, Sod1/3, and Nid2 ranked highest (Fig. 2C). At 6 months, Ly6d, Cdh13, Col12a1, Grn, Psap, and Gsg1l emerged as the top classifiers (Fig. 2D).

Partial correlation network analysis using graphical lasso identified Cdh13 as a central hub, positively associated with Col1a1, Col12a1, Grn, and Spink1 at 1 month and Ly6d at 6 months of age (Fig. 2E-F), suggesting a coordinated biomarker module linked to early matrix expansion and vascular remodeling.

To validate these findings histologically, we performed Picrosirius Red staining for collagen and Cdh13 immunostaining in kidneys from both *Ren1*^*c*^*KO* mice at 1 and 6 months of age and captopril-treated animals at 1, 3, and 6 months of treatment (Fig. 2G). In both models, Cdh13 was markedly upregulated in vessels undergoing concentric hypertrophy and in perivascular inflammatory regions. Collagen deposition increased over time, paralleling disease progression. Control kidneys showed minimal Cdh13 expression, restricted to pericytes, and negligible collagen accumulation. To assess translational relevance, we measured urinary Cdh13 levels by ELISA, confirming its selective detection in *Ren1*^*c*^*KO* mice at both time points (Fig. 2H).

Together, these findings demonstrate that untargeted urinary proteomics, combined with machine learning and multi-model validation, can identify biomarkers that reflect the cellular, structural, and functional trajectory of CAAH. Among these, Cdh13 and collagen isoforms represent promising, non-invasive candidates for monitoring disease activity in the context of chronic RAS inhibition.

### *Ren1*^*c*^*KO* cells develop a matrix-secretory phenotype

To uncover the transcriptomic and biological pathways altered within renin lineage cells during the establishment of CAAH, we performed scRNA-seq in cells sorted as previously described^6^ from mouse kidneys with an intact renin gene (phenotypically WT mice: *Ren1*^*c+/ -*^ ;*Ren1*^*cCre;*^ *R26R*^*mTmG*;^ *Ren1*^*c*^*WT* cells), and from mice with deletion of the renin gene (KO mice: *Ren1*^*c-/-*^;*Ren1*^*cCre;*^ *R26R*^*mTmG*^; *Ren1*^*c*^*KO* cells) at 1 month of age using the 10x *expressed genes between* populations of renin lineage cells including SMCs and JG cells. As anticipated, *Ren1* expression was absent across all cell populations in KO animals (Fig. S2A), but the expression of the surrogate JG cell marker, *Akr1b7*^11^, and the presence of KO cells in integrated clusters enabled the unambiguous identification of these populations (Fig. 3A, Fig. S2B). Within the SMCs, we subclustered these populations to identify four subpopulations representing normal SMCs and three transitional populations expressing both SMC and JG markers (Fig. S2A-F). We characterized the transitional populations as transformed smooth muscle cells (tSMCs) representing recruited renin cells, a process by which renin-lineage derived SMCs reexpress renin under homeostatic threat. Further, putative JG cells formed two distinct clusters, with one cluster displaying an elevated signature of fibrosis (fbJGs) (Fig. S2G).

To identify the gene programs, signaling pathways, and upstream regulators driving these transformed cell states, we first constructed a single-cell trajectory composed of these populations (Fig. 3A). Within each population, we identified the top 20 differentially expressed genes (DEGs) between WT and KO cells (Fig. 3A) and compared them to evaluate which biological pathways were altered in KO mice, which develop CAAH by 1 month of age (Fig. 33B). KO cells became less reliant on oxidative phosphorylation, displayed altered cell adhesion characteristics, underwent epithelial to mesenchymal transition (EMT), increased angiogenesis, and altered KRAS and TNF*α* signaling pathways related to cell growth and proliferation (Fig. 3A-B). Further, KO cells consistently displayed an enriched signature of the senescence-associated secretory phenotype (SASP) (Fig. 3B) that is associated with chronic inflammation and altered local microenvironments as is observed in the pathology of CAAH. In support of this, there is also a uniform enrichment of matrisome-associated genes in all KO cells (Fig. 3B), and enrichment of inflammation-related genes in every KO population except SMCs (Fig. 3B). Additionally, there was a signature of hypoxia present in the fibrotic JG cells (Fig. 3B).

**Fig. 3.**
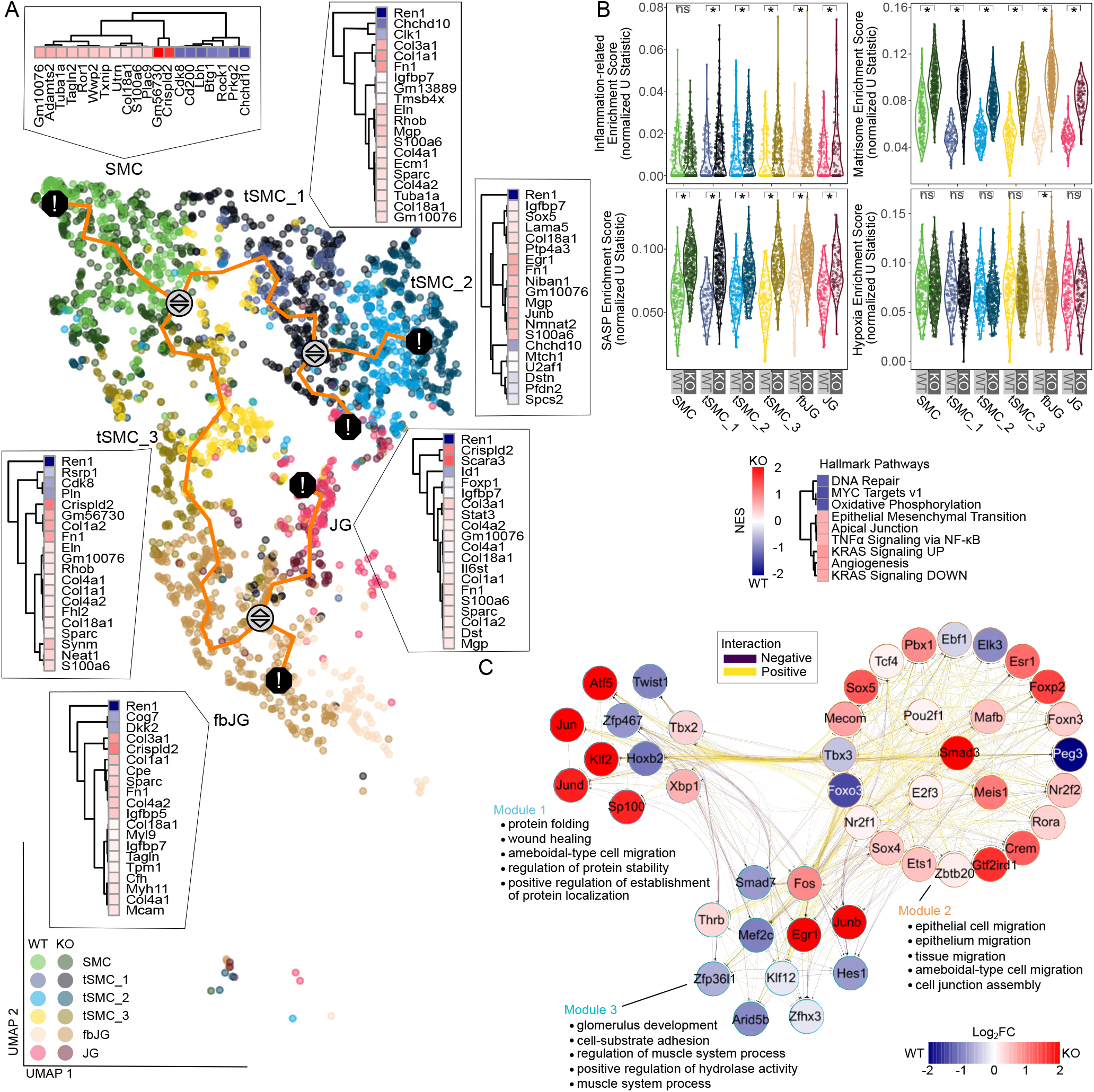
scRNA-seq analysis of Ren1^c+/-^ (phenotypically WT) or Ren1^c-/^- (KO) mice identified enriched signatures of inflammation, fibrosis, extracellular matrix deposition, and a secretory phenotype in KO populations. (A) UMAP visualization of cell populations with inferred trajectory (orange line) showing branch points (gray circles with arrows) and cell fate decisions (black circle with exclamation). For each population, a heatmap depicts the top 20 differentially expressed genes between phenotypically WT and KO mice. (B) Gene signature enrichment scores for inflammation-related genes, the matrisome, the senescence-associated secretory phenotype (SASP), and hypoxia. Pathway enrichment analysis identified loss of oxidative capacity, an epithelial-mesenchymal transition, and inflammation related pathways to be elevated in Ren1 KO cells. (NES=normalized enrichment score. =padj < 0.01. P values are calculated from an estimated marginal means linear model followed by pairwise comparisons between Ren1^c^KO* (KO) and Ren1^c^WT (phenotypically WT) cell populations with the Benjamini-Hochberg multiple testing procedure correction.) (C) Inferred gene regulatory network analysis identified three modules of driver transcription factors composing the renin-lineage trajectory with positive (yellow) and negative (purple) interactions among factors driving the differences observed between trajectory cell populations. For each transcription factor, the color represents the log2 fold change between KO and phenotypically WT cell populations.

We next performed integrated regulatory network analysis across the trajectory to identify putative regulatory transcription factors (TFs) with binding motifs present in the promoter regions of DEGs to infer regulatory relationships. Three distinct regulatory network modules composed the trajectory (Fig. 3C). The most central TFs, those with the most interactions, included *Klf2, Zfp467, Hoxb2, Smad3, Mef2c, Egr1*, and *Klf12* (Fig. 3C). The first TF module, with the central factors *Klf2, Zfp467*, and *Hoxb2*, was associated with gene ontology terms involved in wound healing and protein stability and localization. This may be interpreted to represent the response to changes in blood pressure (BP) homeostasis in which the KO mice are incapable of properly responding, thus contributing to the observed disease physiology with increased extracellular matrix deposition and inflammation. The second TF module, represented by the central TF *Smad3*, was associated with cell migration and cell junctions, corresponding with the accumulation and reorganization of transformed SMCs and JGs observed in CAAH. The third TF module was specifically related to glomerulus development and muscle cell development, again associated with CAAH phenotypic characteristics of arteriolar hypertrophy and the constant, yet ineffective recruitment of more renin null cells.

Together, these findings demonstrate that loss of *Ren1* triggers coordinated transcriptomic reprogramming in renin-lineage cells, promoting fibrotic, proinflammatory, hypoxic, and senescent states that mirror the early pathophysiology of CAAH (Fig. 3B–C). These findings highlight the critical role of renin (and the cells that produce it) in maintaining normal kidney tissue function and the significant consequences of its disruption.

### Chronic RAS inhibition induces inflammation, hypoxia, and a secretory phenotype

To further investigate the underlying mechanisms by which long term RAS inhibition adversely affects the kidneys, we next applied snRNA- and snATAC-seq multiomics to *SMMHC*^*CreERT2*^;*R26R*^*tdTomato*^;*Ren1*^*cYFP*^ mice either untreated or treated with captopril for 1, 3, and 6 months. We applied trajectory analysis to the integrated untreated and captopril-treated populations to compare differences between normal (ctrl) or treated mice. Like *Ren1*^*c*^*KO* mice, treatment with captopril led to the formation of two subpopulations of SMCs representing transformed or transitional cell populations (Fig. 4A). Both populations of these tSMCs represent possible cell fate decisions for cells arising from basal SMCs. Further, the second tSMC population (tSMC 2) represents a transitional state between transformed SMCs and four subpopulations of endothelial cells (Fig. 4A). While evidence of SMCs transdifferentiating directly to endothelial cells is lacking, the reverse is not the case^12,13^, and these populations display clear signatures of endothelial cells (Fig. S3). We also identified the most significantly differentially expressed genes between treated and control cell populations for each cell type along the trajectory (Fig. 4A).

**Fig. 4.**
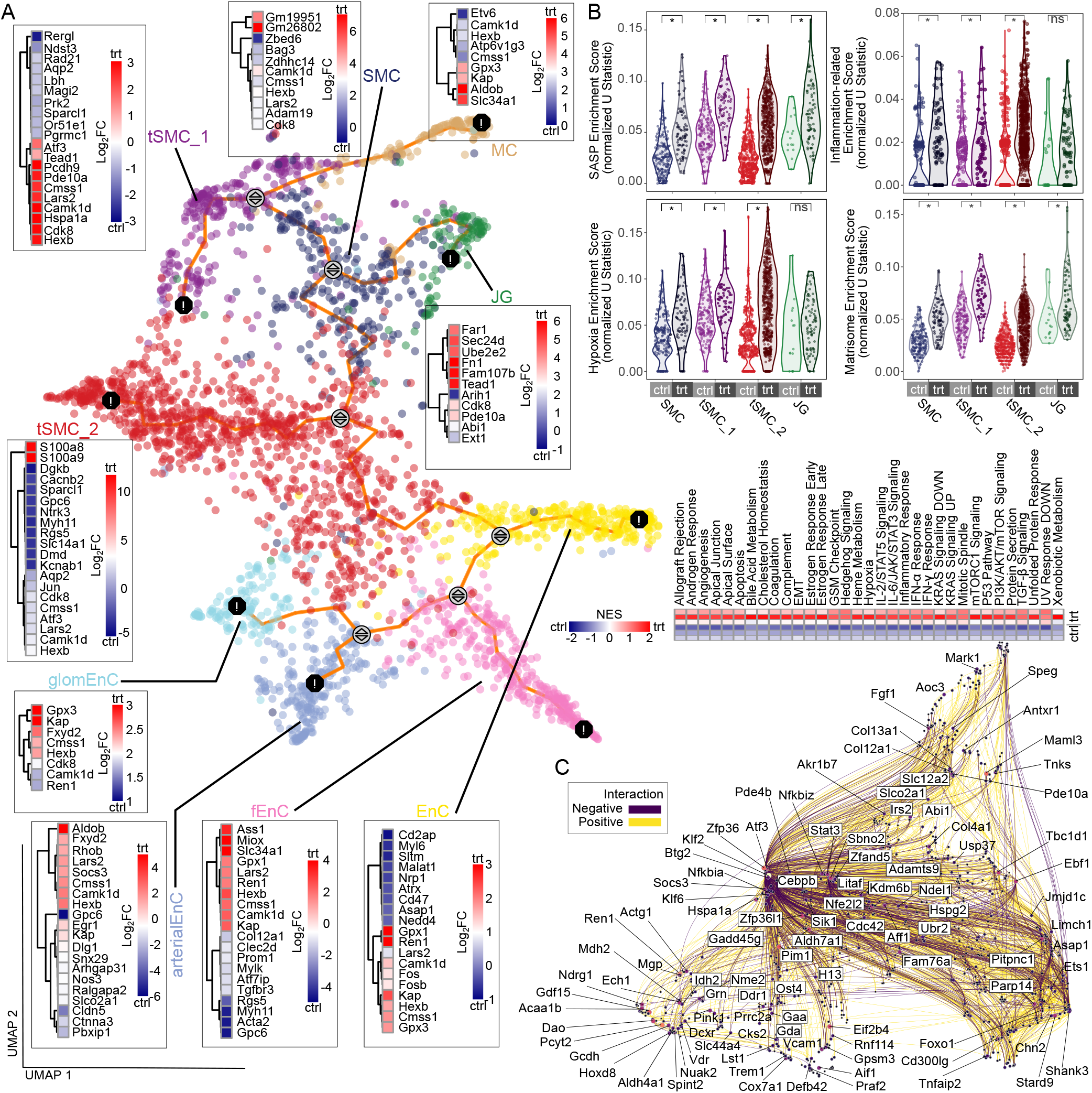
Multiomic analysis of SMMHC^CreERT2^;R26R^tdTomato^;Ren1^cYFP^ mice identifies genes and pathways associated with inflammation, fibrosis, extracellular matrix deposition, hypoxia, and a secretory phenotype in RAS inhibited smooth muscle and renin cell populations. (A)UMAP visualization of cell populations with inferred trajectory (orange line) showing branch points (gray circles with arrows) and cell fate decisions (black circle with exclamation). For each population, a heatmap depicts the top differentially expressed genes between untreated (ctrl) and captopril-treated animals. (B) Gene signature enrichment scores for the senescence-associated secretory phenotype (SASP), inflammation-related genes, hypoxia, and the matrisome. Pathway enrichment analysis identified cell adhesion, epithelial-mesenchymal transition, and inflammatory response pathways to be elevated in RAS inhibited cells. (NES=normalized enrichment score. *=padj < 0.01. P values are calculated from an estimated marginal means linear model followed by pairwise comparisons between treated (trt) and untreated (ctrl) cell populations with the Benjamini-Hochberg multiple testing procedure correction.) (C) Gene regulatory network analysis shows a complex interconnectivity web of genes and positive (yellow) and negative (purple) interactions among genes driving the differences observed between trajectory cell populations.

We next narrowed our focus on investigating the underlying mechanisms that regulate the trajectory between SMCs and JG populations to better understand how long-term RAS inhibition leads to CAAH stemming from these specific cell types in the kidney. To this end, the most significantly elevated hallmark gene pathways in treated populations of SMCs and JGs relative to controls included pathways involved in angiogenesis, cell junctions and adhesion, EMT, inflammatory response signals and pathways, and signaling associated with cell growth and proliferation (Fig. 4B). Again, like *Ren1*^*c*^*KO* mice renin-lineage populations, the senescence-associated secretory phenotype was significantly enriched (Fig. 4B). Further, a gene signature for hypoxia and inflammation was significantly elevated in the treated SMC and transformed SMC populations (Fig. 4B). All treated cell populations from the renin-lineage also had elevated signatures of extracellular matrix deposition and activity (Fig. 4B). These signatures were also present in the other trajectory populations, although to a lesser degree than in the renin-lineage cells known to directly contribute to CAAH (Fig. S3).

We next constructed a gene regulatory network of the SMC, tSMC, and JG cell populations using both transcriptomic and chromatin accessibility data (Fig. 4C). The top 100 genes representing those most central to the network are identified (Fig. 4C). Among these genes are a number related to responses to cellular stress and inflammation (e.g. *Hspa1a, Gdf15, Pim1, Zfp36l1, Sbno2, Nfe2l2, Parp14, Ech1, Cd300lg, Fam76a, Vcam1, Atf3*), mediators of energy metabolism (e.g. *Mdh2, Jmjd1c, Pyct2, Tbc1d1, Acaa1b, Cox7a1, Gcdh, Irs2*), genes involved in cell growth and proliferation (e.g. *Abi1, Stard9, Spint2, Nme2*), or with direct or indirect roles in the regulation of *Ren1* or other RAS system genes (e.g. *Klf2, Foxo1, Fosb, Pitpnc1, Ebf1, Hoxd8, Btg2, Cebpb, Praf2*). Further, significant factors found to be differentially expressed in *Ren1*^*c*^*KO* mice are also present here, including *Klf2, Ebf1*, and *Ets1*, reinforcing that multiple methods of CAAH induction rely on shared pathways driving the observed pathological differences.

### Hypoxia-driven metabolic reprogramming leads to succinate accumulation and NAD^+^ depletion in arteriolar hypertrophy

To evaluate the impact of chronic RAS suppression on renal oxygenation and metabolism, we first studied *Ren1*^*c*^*KO* and control (*Ren1*^*c*^*WT*) mice at 1 and 6 months of age, representing early and advanced disease stages. Hypoxyprobe-1™ staining demonstrated diffuse cortical hypoxia involving glomeruli, tubules, and small arteries (Fig. 5A). Quantitative PCR confirmed a robust hypoxic response, with increased expression of *Hif1α, Hif2α, Epo, Epor*, and *Vegfa* (Fig. 5B). This transcriptional profile was similarly observed in mice treated with captopril for 6 months, supporting the conclusion that both genetic and pharmacological RAS inhibition induce renal hypoxia (Fig. 5C). Untargeted urinary metabolomics (UHPLC-MS/MS) identified 434 nonredundant metabolites across time points. Pathway analysis revealed significant disruption in the TCA cycle and nicotinate/nicotinamide metabolism (Fig. 5D). Notably, urinary succinate levels were consistently elevated in *Ren1*^*c*^*KO* mice at both time points (Fig. 5D). Concurrently, we observed marked depletion of NAD^+^ precursors-niacinamide and 1-methylcotinamidesuggesting impaired NAD^+^ salvage and mitochondrial dysfunction (Fig. 5D). To validate a metabolic shift toward glycolysis, we analyzed renal expression of key glycolytic genes. *Ren1*^*c*^*KO* and captopril-treated kidneys exhibited upregulation of hexokinase-2 (*Hk2*), phosphofructokinase (*Pfkm*), pyruvate kinase-2 (*Pkm*), and lactate dehydrogenase (*Ldha*) confirming Hif-mediated glycolytic reprogramming (Fig. 5E). Immunohistochemistry for Hk2 and Pkm corroborated these findings in both models (Fig. 5F), revealing their distinct localization within renal structures and reinforcing the kidney tissue-level involvement in glycolic reprogramming. In summary, chronic RAS suppression led to renal hypoxia and a Hif-driven metabolic shift marked by succinate accumulation, NAD^+^ depletion, and glycolytic activation (Fig. 5G).

**Fig. 5.**
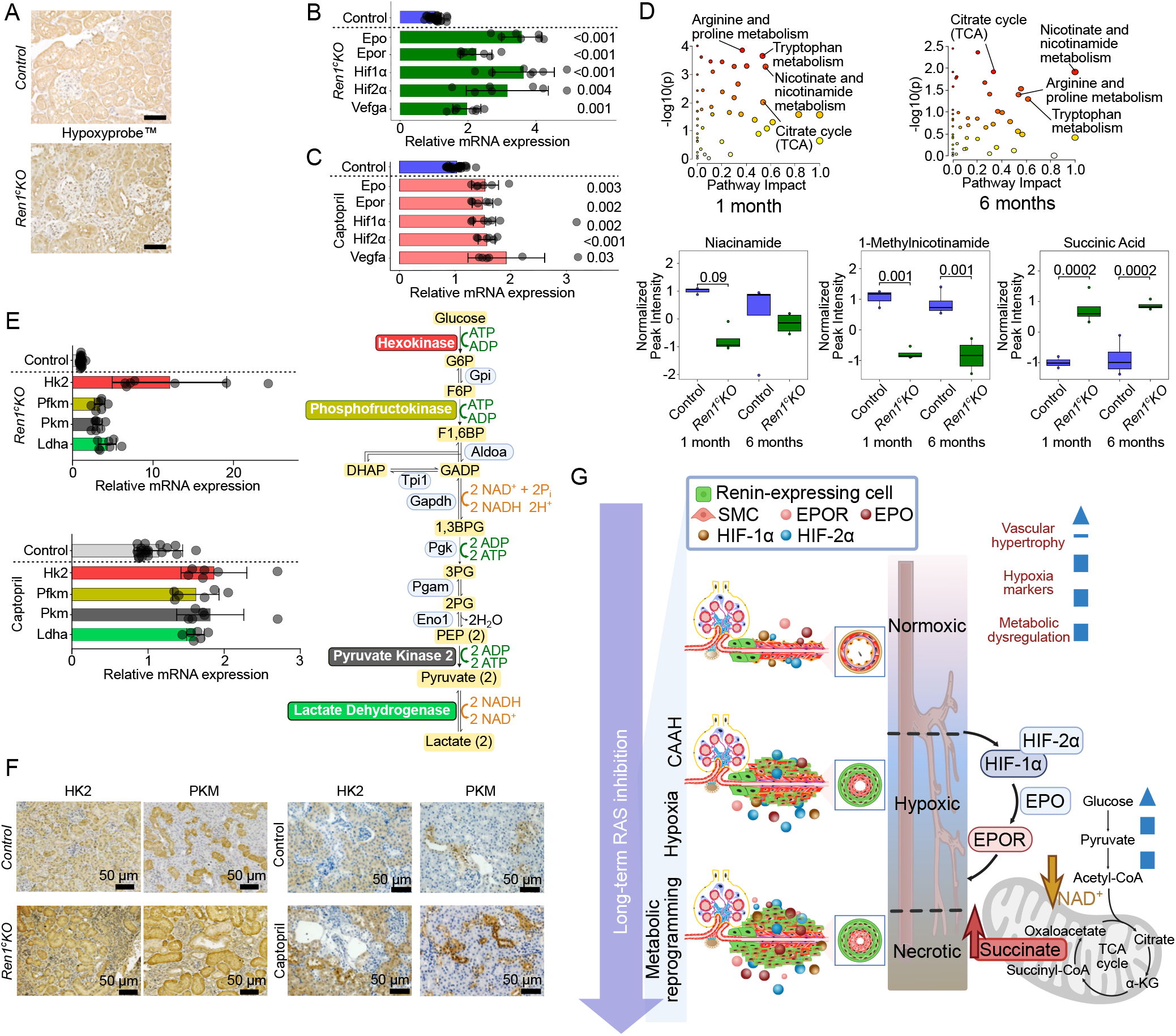
Hypoxia-induced metabolic remodeling in CAAH revealed by urinary biomarkers and gene expression. (A) Hypoxyprobe-1™ immunostaining revealed diffuse cortical hypoxia in glomeruli, tubules, and small arteries in 6-month-old mice. Quantitative PCR demonstrated increased mRNA abundances of hypoxia-responsive genes (Hif1a, Hif2a, Epo, Epor, and Vefga) in 6mo old Ren1^c^KO kidneys (B) and in kidneys from mice treated with captopril (0.5 g/L in drinking water from 2 to 8 months) (C) (p = pairwise t-tests corrected with the Benjamini-Hochberg procedure). (D) Urinary metabolomic pathway analysis highlighted significantly impacted metabolic pathways in Ren1^c^KO mice at 1 and 6 months, with the most prominent being the TCA cycle and nicotinate/nicotinamide metabolism (NAD^+^ depletion and succinate accumulation). Circles represent pathway enrichment (color: p-value, size: impact score). (E) Real time qPCR for hexokinase-2 (Hk2), phosphofructokinase (Pfkm), pyruvate kinase-2 (Pkm), and lactate dehydrogenase (Ldha) in both 6-month-old Ren1^c^KO (upper) and captopril-treated (from 2 to 8 months of age) SMMHC^CreERT2^;R26R^tdTomato;^ Ren1^cYFP^ mice (lower) confirmed Hif-mediated glycolytic reprogramming. All genes are significantly elevated (pairwise T-test with Benjamini-Hochberg multiple testing procedure correction) relative to controls (Ren1cWT or untreated). (F) Immunohistochemistry for glycolytic enzymes HK1, HK2, and PKM demonstrated enhanced expression in renal tissue of 6-month-old Ren1cKO mice. (G) Schematic: Chronic RAS inhibition triggers CAAH and renal hypoxia, activating Hif1α and Hif2α, which drive glycolytic reprogramming, NAD^+^ depletion, and succinate accumulation. Succinate further stabilizes Hifs, sustaining glycolysis, inflammation, and fibrosis, contributing to progressive kidney injury. Abbreviations: CAAH, concentric arteriolar and arterial hypertrophy; Hif, hypoxia-inducible factor; Epo, erythropoietin; Epor, erythropoietin receptor; Vegfa, vascular endothelial growth factor a; NAD^+^ nicotinamide adenine dinucleotide; RAS: renin-angiotensin system.

### Deletion of *Ren1* and chronic RAS pharmacological inhibition identify a CAAH gene signature

Based on our observations of shared pathways, driver genes, and transcription factors between our mouse models, we sought to identify a gene signature of CAAH that effectively discriminates healthy versus diseased kidney samples. We created an initial super set (416 total genes) composed of CAAH-associated genes derived from gene signatures of SASP, inflammation, hypoxia, the matrisome, and our identified protein biomarker genes (Fig. 6A). Next, we ran several thousand permutations of random subsets of 10 genes from the super set and scored both mouse models to identify the set of 10 genes that most discriminated diseased from healthy animals (Fig. 6A). The top scoring set of genes were *Lamp2, Igf1, Cdh13, Col4a2, Sod1, Timp3, Wfdc2, Gpc4, Hsp90ab1*, and *S100a9* (Fig. 6B). Supporting our findings from the proteomics analysis, Lamp2, Cdh13, Sod1, Wfdc2 were identified both here and in the urine as possible biomarkers. Further, while Col4a2 was not identified in the urine, two other collagens (Col12a1 and Col1a1) were, and Col4a2 is a major structural component of basement membranes and its signature in CAAH disease models is arguably a hallmark of the observed loss of GFR.

**Fig. 6.**
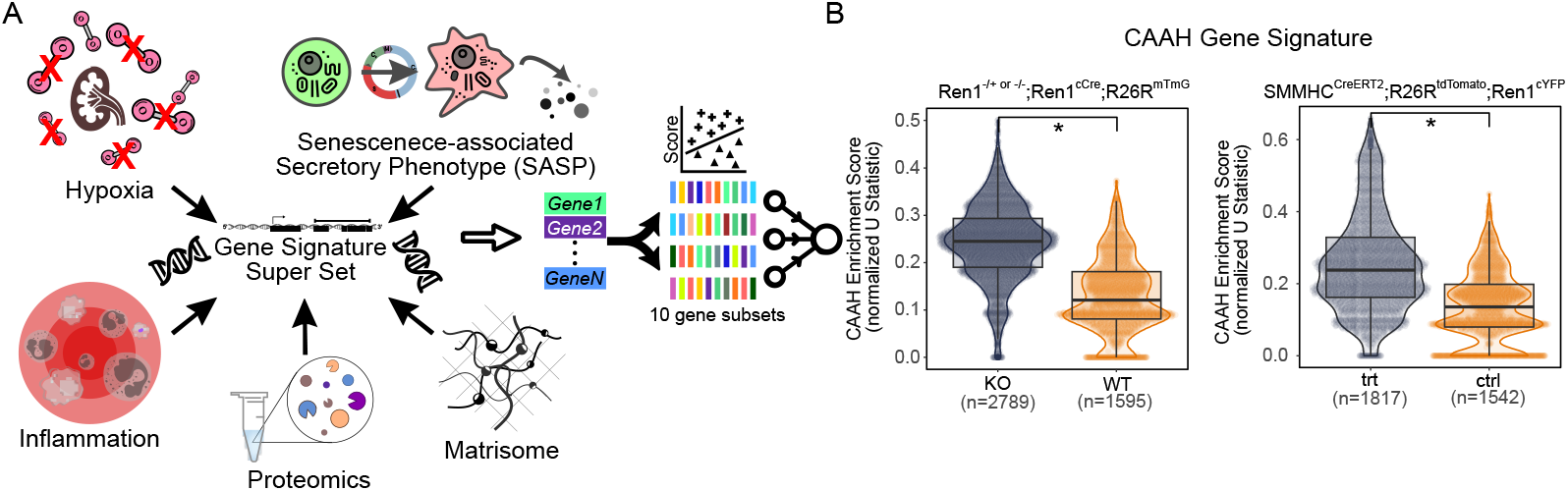
Ten-gene signature discriminates between RAS inhibited and Ren1 genetic ablataion models compared to matched controls. (A) Conceptual cartoon of the identification of a CAAH gene signature. A superset was constructed composing 416 genes from signatures of hypoxia, the senescence-associated secretory phenotype, inflammation, the matrisome, and putative biomarker proteins from our proteomic analysis. Several thousand permutations of randomly selected subsets of 10 genes were scored in both RAS inhibited and Ren1^c^KO cells and the gene set with the greatest differential score between untreated/wildtype and treated/knockout animals was selected. (B) The 10 gene CAAH gene signature was scored in all cells from SMMHC^CreERT2^;R26R^tdTomato;^ Ren1^cYFP^ and Ren1^c-/-^ or Ren1^c+/^ -;Ren1^cCre^;R26R^mTmG^ mice and showed significant differences (Welch’s T: p<0.01) between conditions in both models.

## Discussion

Using integrated single-cell transcriptomics, proteomics, metabolomics, and advanced imaging, we uncovered the molecular and cellular mechanisms underlying CAAH—a silent, progressive renal vasculopathy induced by chronic RAS suppression. In *Ren1*^*c*^*KO* mice, loss of angiotensin II signaling led to renal dysfunction characterized by reduced GFR and elevated BUN. These functional impairments were preceded by striking structural abnormalities, including concentric arteriolar thickening, perivascular immune infiltration, interstitial fibrosis, and tubular enlargement. These findings confirm that renin-lineage cells, beyond their endocrine function, are essential for maintaining mural cell identity and renal vascular integrity^14,15^.

Single-cell transcriptomic analysis revealed that reninlineage cells lacking renin expression (*Renin*^*null*^/*Ren1*^*c*^*KO* cells) undergo dramatic transcriptional reprogramming, acquiring stress-responsive and profibrotic profiles. Notably, these cells expressed *Sparc*, a matricellular protein that modulates ECM synthesis^16^, and *Egr1*, a zincfinger transcription factor implicated in vascular remodeling and TGF-*β* signaling^17,18^. The upregulation of *Egr1* was accompanied by increased expression of its targets including *Col1a1*^19,20^, supporting its role as a central driver of fibrosis. Renin-lineage cells also produced multiple ECM components (*Col1a1, Col3a1, Col4a1, Fn1*), whose accumulation likely contributes to vascular narrowing and downstream ischemic injury. The expression of *Scara3*, an oxidative scavenger, suggests an adaptive response to local hypoxia^21^. These changes illuminate how renin-lineage cell plasticity actively contributes to structural and functional decline.

To uncover translational biomarkers of early disease, we employed untargeted urinary proteomics. Col1a1 was detectable exclusively in *Ren1*^*c*^*KO* urine as early as one month—prior to overt functional impairment—suggesting early fibrotic remodeling. Histological validation with Picrosirius red staining confirmed progressive collagen deposition in renal tissues, offering a potential noninvasive marker of early ECM expansion and CAAH establishment through type-I collagen-targeted in vivo imaging^22,23^. Similarly, Cdh13, encoding T-cadherin, was elevated in both renal tissue and urine at 1 and 6 months of age. Its role in vascular remodeling and association with proliferative vascular disorders^24–27^ positions Cdh13 as a compelling biomarker candidate. These findings reinforce the potential for urinary ECM and vascular remodeling proteins to signal early CAAH establishment.

Crucially, these pathological features were also observed in mice treated with captopril, a clinically used RAS inhibitor. Prolonged pharmacologic RAS suppression recapitulated structural vascular injury, including concentric arteriolar thickening, underscoring the clinical relevance of our findings. Notably, vascular remodeling preceded measurable renal functional decline, highlighting the kidney’s remarkable compensatory capacity and reinforcing the need for early detection strategies.

We uncovered a mechanistic axis linking chronic RAS suppression—whether genetic or pharmacologic—to renal hypoxia and metabolic reprogramming. Hypoxyprobe-1^™^ staining revealed widespread cortical hypoxia in both *Ren1*^*c*^*KO* and captopril-treated mice. These changes were accompanied by transcriptional activation of *Hif1α, Hif2α*, and downstream targets such as *Epo* and *Epor*, indicating robust Hif pathway activation. Urinary metabolomics revealed elevated succinate—a TCA intermediate that stabilizes Hif1*α* by inhibiting prolyl hydroxylases—suggesting a feed-forward loop of hypoxia and metabolic stress. Simultaneously, depletion of NAD^+^ precursors (niacinamide, 1-methylnicotinamide) pointed to impaired mitochondrial redox balance. Consistent with Hif-mediated metabolic shifts, glycolytic enzymes (*Hk2, Pfkm, Pkm*, and *Ldha*) were upregulated in renal tissue, and protein localization studies confirmed glycolytic reprogramming across renal compartments. Together, these findings illustrate a convergent pathway of vascular remodeling, hypoxia, and energy failure that drives disease progression in both models of RAS suppression.

Beyond proteomics and metabolomics, we identified a robust transcriptional signature of CAAH composed of *Igf1, Timp3, Gpc4, Hsp90ab1, S100a9, Lamp2, Cdh13, Sod1, Wfdc2*, and *Col4a2*. These markers converge on key biological processes including vascular remodeling, fibrosis, inflammation, oxidative stress, and autophagy. For instance, Igf1 promotes renal fibrosis and hemodynamic stress^28–31^; Timp3 mitigates ECM turnover and fibrosis^32–34^; Gpc4 and Hsp90ab1 influence podocyte injury and vascular stress^35,36^; S100a9 acts as a DAMP molecule exacerbating renal inflammation^37–39^; Lamp2 safeguards autophagic flux under stress^40–42^; Sod1 counters oxidative injury^43,44^; Wfdc2 mediates fibrotic responses^45–48^; and Col4a2 supports hypoxia-driven matrix remodeling^49,50^.

Importantly, several of these pathways present tractable targets for therapeutic intervention. Agents such as Teprotumumab (IGF-1R inhibition), Paquinimod (S100A9 blockade), Ilomastat (broad-spectrum MMP inhibition impacting Timp3 and Gpc4 pathways), and prolyl hydroxylase (PHD) inhibitors (targeting Hif1*α*-mediated *Col4a2* regulation) offer near-term translational opportunities. Further, NAD^+^ supplementation is reported to be protective in chronic kidney disease models^51^, and multiple avenues exist to target succinate accumulation either through the inhibition of succinate dehydrogenase using dimethyl malonate or supplementation with itaconate^52,53^, for example, or the targeting of the succinate receptor^54^. These compounds, many already in clinical trials or approved for other indications, facilitate potential repurposing strategies aimed at preventing or mitigating CAAH.

Collectively, our findings uncover a shared pathological program initiated by both genetic and pharmacologic suppression of angiotensin II signaling. This program is defined by mural cell reprogramming, vascular fibrosis, renal hypoxia, metabolic stress, and eventual tubular injury. Crucially, these changes occur early and silently, before loss of renal function becomes clinically apparent. The identification of candidate urinary biomarkers— collagens (Col1a1, Col12a1), Cdh13, succinate, and niacinamide — opens the door to early detection and monitoring strategies for patients on chronic RAS inhibition. Future studies should prioritize validation of these biomarkers in human cohorts and explore therapeutic strategies aimed at restoring metabolic balance or disrupting Hif- and ECM-driven injury cascades.

## Extended Methods

### Renin knockout model of CAAH

Ren1c null mice were generated as previously described^55^. To generate *Ren1*^*c-/-*^;*Ren1*^*cCre*^;*R26R*^*mTmG*^ mice, we crossed *Ren1*^*c-/-*^ mice with *Ren1*^*cCre*^ mice^56^ and *R26R*^*mTmG*57^. In this mouse model, renin lineage cells are renin null and GFP+. *Ren1cKO* mice from both sexes were studied at 1, 3, and 6 months representing progressive and worsening stages of the arterial disease and compared to their respective controls.

### Pharmacological RAS inhibitor model of CAAH

We developed a mouse model to study the transformation of smooth muscle cells (SMCs) as the animals developed CAAH while receiving a RAS inhibitor. *SMMHC*^*CreERT2*^;*R26R*^*tdTomato*^;*Ren1*^*cYFP*^ mice track cells derived from the SMC lineage (tdTomato+) after tamoxifen administration (100 mg/kg, intraperitoneal for 3 consecutive days). Two-month-old male (the Myh11-Cre^ERT2^ transgene is located on the Y chromosome) *SMMHC*^*CreERT2*^;*R26R*^*tdTomato*^;*Ren1*^*cYFP*^ mice received captopril (100mg/kg/day) in their drinking water *ad libitum* for 1, 3, and 6 months (n=6 per timepoint) and were compared to control mice (n=6 per timepoint) with unadulterated drinking water for the same lengths of time. In this model VSMCs are labeled red (tdTomato+) upon tamoxifen injection whereas cells currently expressing renin are labeled yellow (YFP+)^58^.

All animals were housed in the University of Virginia’s vivarium facilities equipped with controlled temperature and humidity conditions under a 12 hours dark/light cycle. All protocols were performed per the Guidelines for the Care and Use of Laboratory Animals published by the United States National Institutes of Health (https://grants.nih.gov/grants/olaw/guide-for-the-care-and-use-of-laboratory-animals.pdf) and approved by the University of Virginia Animal Care and Use Committee. Adult mice were killed after anesthesia with tribromoethanol (intraperitoenal, 300 mg/kg or isoflurane) followed by cervical dislocation. Kidneys were processed for histological, 3D, immunohistochemical and multiomics analyses as indicated below.

### Blood pressure measurement using radiotelemetry

Blood pressure was measured using a radiotelemetry device (Data Sciences International, Orange, CA). During implantation of the radio-telemeters, mice were fully anesthetized with isoflurane and buprenorphine was used as an analgesic (0.06mg/mL). Radio-telemeters (HD-X10 or PA-C10) were implanted in a subcutaneous pocket on the right side of the mouse’s body. Catheters were implanted into the left carotid artery with the probe extending to the aortic arch. Following surgery and recovery (7 days), heart rate, systolic blood pressure, diastolic blood pressure, and mean arterial pressure were recorded and processed for an additional 7 days using Dataquest A.R.T 20 software (DSI).

### Transcutaneous measurement of GFR by FITC-sinistrin

A noninvasive clearance device (NIC-Kidney Device; MediBeacon, Mannheim, German) was used to measure glomerular filtration rate (GFR). Briefly, a small fluorescence detector was fixed on the depilated back of isoflurane-narcotized (isoflurane 4%; 0.5-1.0 L/min, O2) mice using a double-sided adhesive patch. After beginning fluorescent measurements, FITC-sinistrin (10 mg/100 g body weight dissolved in 0.9% NaCl) was injected intravenously and the elimination of FITC-sinistrin was measured transcutaneously in conscious mice for 60-90 minutes^59^.

### RNA Isolation and Real-Time RT-PCR Analysis

Quantitative real-time PCR (qPCR) was conducted on kidney cortex samples to assess gene expression. Total RNA was extracted using the TRIzol reagent (Thermo Fisher Scientific) and the RNeasy Mini Kit (Qiagen, Germantown, MD). Reverse transcription was performed using oligo(dT) primers and M-MLV reverse transcriptase (Promega, Madison, WI) at 42°C for 1 hour, following the manufacturer’s protocol. qPCR was performed using SYBR Green I (Thermo Fisher Scientific) on a CFX Connect system (Bio-Rad Laboratories, Hercules, CA). The following primers were used: *Hif1α*, forward: 5’-TGACGGCGACATGGTTTACA-3’, reverse: 5’-ACTGGGCCATTTCTGTGTGT-3’; *Hif2α*, forward: 5’-GTCGAGGAAGGAGAAATCCCG-3’, reverse: 5’-GTTATCCATTTGCTGGTCGGC-3’; *Epo*, forward: 5’-CCTCATCTGCGACAGTCGAGTTC-3’, reverse: 5’-CAGTCTGGGACGTTCTGCACAAC-3’; *Epor*, forward: 5’-GAGACCCTCCCAAGGAACA-3’; reverse: 5’-CTCCGACAGACTGACTCGC-3’; *Vegfa*, forward: 5’-TATTCAGCGGACTCACCAGC-3’, reverse: 5’-CCTCCTCAAACCGTTGGCA-3’;*Hk2*, forward: 5’-GGAACCGCCTAGAAATCTCC-3’; reverse: 5’-GGAGCTCAACCAAAACCAAG-3’; *Pfkm*, forward: 5’-TTACTCAGTGGAACACCGCC-3’; reverse: 5’-GTTTATCCCCCGATTCAGGT-3’; *Pkm*, forward: 5’-GTCTGAATGAAGGCAGTCCC-3’; reverse: 5’-GTCCGCTCTAGGTATCGCAG-3’; *Ldha*, forward: 5’-GTGCCCAGTTCTGGGTTAAG-3’; reverse: 5’-CTGGGTCCTGGGAGAACAT-3’ and *Rps14* (housekeeping gene): Forward: 5CAGGACCAAGACCCCTGGA-3’ Reverse: 5’-ATCTTCATCCCAGAGCGAGC-3’.

Expression levels of *Hif1α, Hif2α, Epo, Epor, Vegfa, Hk2, Pfkm, Pkm*, and *Ldha* mRNA were normalized to *Rps14*, and relative expression changes were quantified using the ΔΔCt method, with results expressed as fold changes relative to control mice.

### Blood chemistry

Animals were anesthetized with tribromoethanol (300 mg/kg). Blood was collected by cardiac puncture and placed into tubes containing EDTA or heparinized plasma separator tubes (BD Microtainer, Becton Dickinson). Basic metabolic panel analysis was performed by the University of Virginia Hospital clinical laboratory^7^.

### ELISA for renin in plasma

Plasma specimens were obtained from blood after centrifugation at 1000*×sss* g at 4 °C for 20 minutes. Renin concentration was determined using ELISA (Enzyme-Linked Immunosorbent Assay) following the manufacturer’s instructions (RayBiotech, Norcross, GA)^7^.

### Quantification of Cadherin 13 in urine samples

Cadherin 13 quantification was performed using ELISA commercial kits (Invitrogen, Carlsbad, CA). The median values of Cadherin 13 in the urine samples were expressed in mg/g of creatinine.

### Histological and immunohistochemical analyses

Mice were anesthetized with tribromoethanol intraperitoneally (300 mg/kg). Kidneys were removed, fixed overnight in Bouin’s solution at room temperature (RT) or 4% paraformaldehyde (PFA) or formalin at 4^*°*^C, and embedded in paraffin. Immersion fixation as opposed to perfusion fixation was performed so that RNA extraction for gene expression analysis could be performed for the same animal. Sections (5 *µ*m) from Bouin’s-fixed, paraffin-embedded kidneys were stained with Hematoxylin and Eosin (MilliporeSigma, Burlington, MA) to examine kidney morphology and picrosirius red stains to study collagen networks (Polysciences, Warrington, PA). Immunostaining was performed as previously described^60^. Briefly, 5 *µ*m sections of Bouin’s-fixed, paraffin-embedded kidneys were incubated overnight at 4°Cwith a rabbit polyclonal anti-mouse renin antibody (1:500) generated in our laboratory^60^, mouse anti *α*-SMA monoclonal antibody (1:10,000; MilliporeSigma), hexokinase II rabbit monoclonal antibody (1:200; Cell Signaling), PKM2 rabbit monoclonal antibody (1:800; Cell Signaling), or mouse/human anti-cadherin 13 polyclonal antibody (1:200, Invitrogen, Waltham, MA) and biotinylated secondary goat anti–rabbit IgG or horse anti–mouse IgG (1:200; Vector Laboratories, Newark, CA) for cadherin 13 and renin, or *α*-SMA respectively. Staining was amplified using the Vectastain ABC kit (Vector Laboratories) and developed with 3,3-diaminobenzidine (MilliporeSigma). The sections were counterstained with hematoxylin (MilliporeSigma), dehydrated, and mounted with Cytoseal XYL (Thermo Fisher Scientific, Waltham, MA).

### Detection of Tissue Hypoxia

Tissue hypoxia was detected using the Hypoxyprobe1^™^ Plus Kit (Hypoxyprobe, Inc., catalog no. HP2-100Kit), which utilizes pimonidazole hydrochloride, a 2-nitroimidazole compound that forms stable adducts with thiol-containing proteins in hypoxic cells (pO_2_ *<*10 mmHg). Mice were injected intraperitoneally with pimonidazole hydrochloride at a dose of 60 mg per kg body weight, diluted in sterile 0.9% saline. After 90 minutes of circulation, mice were euthanized under deep anesthesia, and kidneys were perfused with cold PBS, harvested, and fixed in 4% paraformaldehyde overnight at 4^*°*^C, followed by paraffin embedding.

Five-micrometer paraffin sections were deparaffinized, rehydrated through graded alcohols, and subjected to heat-induced epitope retrieval using 10 mM citrate buffer (pH 6.0) for 10 minutes. Endogenous peroxidase activity was quenched with 3% hydrogen peroxide, and sections were blocked in 5% normal rabbit serum for 1 hour at room temperature. Slides were incubated overnight at 4^*°*^*C*with the anti-pimonidazole, FITC-conjugated IgG rat or mouse monoclonal antibody (provided in the kit, clone 11.23.22. R; 1:100). After washing, sections were incubated with a 1:100 dilution of peroxidase conjugated anti-FITC secondary reagent and developed with DAB substrate.

Sections were counterstained with hematoxylin, dehydrated, cleared, and mounted using Permount^™^.

### Isolation of single cells

To isolate single cells, first the kidneys were excised and decapsulated. Then, the kidney cortices were dissected, minced with a razor blade, and transferred into a VIA Extractor™ tissue pouch (CytivaTM) with 5 mL of enzymatic solution (0.3% collagenase A [MilliporeSigma], 0.25% trypsin [MilliporeSigma], and 0.0021% DNase I [Millipore-Sigma]). The pouches were placed flat inside a VIA Extractor^™^ tissue disaggregator (CytivaTM) (200 rpm) for 12 minutes at 37^*°*^C. The tissue was collected and centrifuged at 800*×* g for 4 minutes at 4^*°*^*C*using a Sorvall RT7 refrigerated centrifuge (Sorvall). The cell pellet was resuspended in fresh buffer 1 (130 mM NaCl, 5 mM KCl, 2 mM CaCl2, 10 mM glucose, 20 mM sucrose, 10 mM HEPES, pH 7.4), and the suspension was poured through a sterile 100 *µ*m nylon cell strainer (Corning Inc.) and washed with buffer 1. The flow-through was poured through a sterile 40 *µ*m nylon cell strainer (Corning Inc.) and washed with buffer 1. The flow-through was centrifuged at 800*×* g for 4 minutes at 4^*°*^*C*using a Sorvall RT7 refrigerated centrifuge (Sorvall). Residual red blood cells were lysed by Red Blood Cell Lysis Buffer (Sigma-Aldrich). The cell pellet was resuspended in 1.5 mL of resuspension buffer: PBS, 1% FBS, 1 mM EDTA, DNase I (MilliporeSigma). The dead cells were labeled with DAPI (MilliporeSigma). Cells were analyzed and sorted using an Influx Cell Sorter (Becton Dickinson) at the Flow Cytometry Core Facility of the University of Virginia. Finally, cells were collected in DMEM with 20% FBS and used for single-cell capture and downstream applications^6^.

#### Single-cell (sc) capture and RNA sequencing

Freshly isolated FACS sorted kidney cells were resuspended in dPBS containing 0.04% BSA and the volume adjusted to yield 1000 cells/*µ*L. Single cells were loaded and captured using the Chromium Controller System (10X Genomics, Pleasanton, CA) following the manufacturer’s recommendation^61^ for the Chromium Next GEM Chip G with reagents of Chromium Next GEM Single Cell 3^*′*^ Reagent Kits v3.1 Dual Index (10X Genomics). Finally, the completed libraries were sent to MedGenome (Foster City, CA) for sequencing using a 28-90 paired-end configuration.

#### Single-cell multiomic RNA and ATAC sequencing

Single cell multiome libraries were constructed from freshly isolated FACS sorted kidney cells resuspended in dPBS with 0.04% BSA at a concentration of 1000 cells/*µ*L. Single cells were processed using the Chromium Controller System (10X Genomics, Pleasanton, CA) following the manufacturer’s recommendation^61^ for the 10x Genomics Chromium Next GEM Chip J with reagents of the Chromium Single Cell Multiome ATAC + Gene Expression kit. Completed libraries were sent to MedGenome (Foster City, CA) for sequencing using a 28-90 paired-end configuration for scRNA multiome libraries and a 50-49 paired-end configuration for scATAC multiome libraries.

### scRNA-seq and scMultiome analyses

#### Alignment and feature-barcode matrix generation

Fastq files from the 10X Genomics Single Cell Gene Expression platform were processed using the Cell Ranger pipeline (version: `cellranger-7.2.0`) and the `refdata-gex-mm10-2020-A` reference data. This pipeline trims non-template sequence and aligns reads using the STAR aligner^62^. Following read mapping, reads were grouped by barcode, UMI, and gene with reads mapped to genes calculated using the UMI-based counts. Filtered UMIs were linked to barcodes with the feature-barcode matrices used for downstream analysis. Fastq files from the Single Cell Multiome ATAC + Gene Expression libraries were intially processed using the Cell Ranger ARC pipeline (version: `cellranger-arc-2.0.2`) and the `refdata-cellranger-arc-mm10-2020-A-2.0.0` reference data. As with Single Cell Gene Expression libraries, the filtered feature-barcode matrices were used for further analysis.

#### Sequencing quality control

Feature-barcode matrices produced by Cell Ranger were analyzed in R using Seurat v. 5.2.1^63,64^. For scRNA samples, cells with mitochondrial content below the 92.5^^^th percentile or hemoglobin content less than the 97.5^^^th percentile were retained for downstream analysis. Cells were also required to contain a number of identified RNA features between the 2.5^^^th and 97.5^^^th percentiles, as extreme values are indicators of low quality or doublet cells^65,66^.

#### Cell clustering and annotation

*scRNA-seq (*Ren1^^^c+/- or -/-^;Ren1^cCre^;R26R^mTmG^^^ *mice)*. To address cell-cell variability due to batch and cluster data, we utilized standard integration workflows in Seurat v5^64^ following normalization using the functions `NormalizeData(), FindVariableFeatures(), ScaleData(), RunRCA(), and IntegrateLayers()` using Harmony (v. 1.2.0) integration. For initial clustering and annotation, cell clusters were generated using the `FindNeighbors()` function on the top 1:15 dimensions and `FindClusters()` with a resolution of 1.5 and using algorithm 1. Uniform Manifold Approximation and Projection (UMAP) was generated on the harmony reduction with the following settings: `n.neighbors = 15L, n.epochs = 500, min.dist=0.3, spread=0.5, seed.use=99`. The top 25 differentially expressed genes present in at least 1% of the cells in a cluster and with a minimum 1 log fold change between clusters (see Seurat `FindAllMarkers`) were defined as markers for each cluster. To perform cluster annotation, an iterative approach was performed by comparing the top markers to previously identified markers from mouse cell atlases and a commercial database (cellKb^67^). First, markers from internal data were analyzed for expression in each identified clusters. Second, marker expression was compared with cell identity markers from three atlases of developing mouse kidney cells^68–70^. Third, the identified markers were matched with cell signatures using the commercial CellKb database to create rank ordered lists of top matching gene signatures to cell signatures. The resulting top ranked hits from each approach were analyzed together and cell cluster identities manually annotated.

After identifying renin-lineage cell populations (SMCs, tSMCs1-3, fbJG cells, and JG cells), we then subsetted the Seurat object on just these cell populations as they represent the known cell types that contribute to CAAH. We reperformed neighbor analysis `(FindNeighbors())` using canonical correlation analysis (CCA) and found clusters `(FindClusters())` with the following parameters: `resolution=0.75, random.seed=99, algorithm=1`. The UMAP of the clusters found following CCA reduction was produced using `RunUMAP()` with the following parameters: `seed.use=99, n.neighbors = 15L, n.epochs = 500, min.dist=0.01, spread=1, local.connectivity = 1.5L, repulsion.strength = 1, negative.sample.rate = 25L`. Cluster annotations were refined for the subclustered renin-lineage cell populations as described above. Subsequent analyses used these refined renin-lineage cell annotations.

*scMultiome (*SMMHC^CreERT2^;R26R^tdTomato^;Ren1^cYFP^ *mice)*. Joint chromatin accessibility and gene expression (scATAC- and scRNA-seq) multiomic data was analyzed using Seurat v5^64^ and Signac v^71^. scATAC-seq data was normalized using term-frequency inverse-document-frequency (see `RunTFIDF()`) and reduced using latent semantic indexing (LSI) (see `RunSVD()`). The scRNA-seq component was normalized and variance-stabilized using a regularized negative binomial regression implemented in R (see Seurat `SCTransform()`), while controlling for the cell cycle and mitochondrial and hemoglobin percentage^63,72^. scRNA-seq was reduced using `RunPCA()`. Weighted nearest neighbor analysis (parameters: `reduction.list = list(“pca”, “lsi”), dims.list = list(1:50, 2:50)`) was performed using `FindMultiModalNeighbors()` to identify the initial shared clusters. scRNA-seq data was then integrated across sample layers using `IntegrateLayers(`) and the fast mutual nearest neighbors (FastMNN) dimensionality reduction^73^. Marker genes for each cluster and cell type annotation was performed as described in the scRNA-seq data for *Ren1c* mice.

The top transcription factor regulators for each cell population was calculated using two approaches. First, the combined expression and motif accesibility scores were calculated using the R package Presto^74^. Second, the chromVAR^75^ deviation scores were calculated for known motifs in each cell cluster. Top scoring transcription factor motifs were identified for each cluster using `FindAllMarkers()` for the `chromvar` assay using the following parameters: `random.seed=99, min.pct=0.01, logfc.threshold=1`. Potential regulators required an adjusted p-value less than 0.01, an absolute log fold change greater than 0, and were ranked according to the averaged RNA and motif accessibility area under the curve (AUC).

#### Trajectory analysis

For *Ren1*^*c+/-*^ (WT) or *Ren1*^*^*^*c-/ -*^*^*^ (KO) mice we learned the trajectory of renin-lineage cells using the `learn graph()` function from monocle3^76–78^. The trajectory was then visualized on the UMAP and we defined the basal smooth muscle cell (SMC) population as the root node. For *SMMHC*^*CreERT2*^;*R26R*^*tdTomato*^;*Ren1*^*cYFP*^ mice treated with captopril or untreated, we performed the same procedure including both renin-lineage cells and glomerulus associated endothelial cell populations. We visualized the resulting trajectory (via `learn graph()`) on the combined UMAP and set the SMC population as the root node. We next identified genes that changed as a function of pseudotime (see `graph test() in monocle3`) and kept those with a q-value less than 0.05.

#### Differential gene expression analysis

Differentially expressed genes (DEGs) within populations between either *Ren1cKO* and *Ren1cWT* or captopril-treated and untreated *SMMHC*^*CreERT2*^;*R26R*^*tdTomato*^;*Ren1*^*cYFP*^ mice was calculated using the `FindMarkers()` function from Seurat using the Model-based Analysis of Single-cell Transcriptomics approach, MAST^79^, with default parameters. The top DEGs, defined as those significantly differently expressed genes (Bonferroni adjusted p-value *<* 0.01) with the greatest log2 fold change, were plotted as heatmaps using the `pheatmap()` function from the `pheatmap R` package^80^ for each comparison.

#### Gene regulatory network analysis

*scRNA-seq*. To construct an inferred gene regulatory network (iGRN), we began with the renin-lineage only `Seurat` object composing the trajectory defined above. To identify genes that changed as a function of cell type, we used the `fit models()` function from monocle3^76–78^, then selected only those genes with a q-value less than 0.01, that were expressed in at least 1 cell, and we excluded the intercept terms. We then added the `monocle3` pseudotime values back to the original `Seurat` object using the `AddMetaData()` function. To identify candidate transcription factors (TFs), we used a list of TFs with known motifs from the `Tranfac201803 Mm MotifTFsF` database provided in the `IReNA R` package^81^. Next, we retained only the known TFs that were present in the `Seurat` renin-lineage object and expressed in at least 5% of cells. We then identified genes that changed as a function of pseudotime using the get SmoothByBin PseudotimeExp()^81^ function with a bin size of 50 and a fold change threshold of Q95. From the smoothed expression profile, only genes with a fold change greater than 0.25 were retained for further analysis. To identify the optimum number of TF clusters, we used the most common optimum cluster number using the elbow, silhouette, and gap methods implemented via the `fviz_nbclust()` function in the `factoextra v.1.0.7` package^82^. Next, we performed K-means clutering using the `clustering_Kmeans()` function in `IReNA`^81^. We added Ensembl gene ids to each gene using `add_ENSID()` and inferred possible regulatory relationships for each gene in the filtered expression profile via the `GENIE3` algorithm^83,84^. To refine these relationships, the promoters of each gene were then scanned to determine whether binding motifs of the predicted transcription factors were present using the `identify_region_tfs()` function and the `motifmatchr R package`^85^. Next, we constructed the regulatory network via the `network analysis()` function. We performed gene ontology analysis for the identified regulatory modules using `enrich_module()`^81^. Finally, the resulting network was plotted using `Cytoscape`^86^.

*scMultiome*. To construct a gene regulatory network (GRN), we included only the renin-lineage cell populations along the identified trajectory as described above. We then created the intial regulatory network using the `initiate grn()` function from the R package `Pando v.1.1.1`^87^. We chose candidate regions derived from the `LinkPeaks()` function in `Seurat`^64^. Next, we identified the presence of motifs in the candiate regions using the`find_motifs()` function from `Pando` using the `Pando` curated motif collection. Next, using the set of genes that changed as a function of pseudotime described above, we first identified which of those genes were differentially expressed between captopril-treated and untreated cells using `FindMarkers()` and the `MAST test`^79^ with default parameters. Tbose genes with an adjusted p-value less than 0.01 were then provided to a modified `Pando infer_grn()` function to enable the `GREAT`^88,89^ peak to gene method using the mouse genome. The resulting GRN was then visualized using `get_network_graph()`and `plot_network_graph()` from `Pando`^87^.

#### Gene set enrichment and pathway analysis

We evaluated the enrichment of annotated gene sets from the Molecular Signatures Database^90,91^ for mouse hallmark, regulatory target, curated, cell signature, biological process, cellular component, and molecular function gene sets. For each gene set, we scored individual cells using the Single Pathway analysis in Single Cells (`SiPSiC v.1.4.3`) R package^92^. All pathway scores were then z-score normalized and plotted as heatmaps using the R package pheatmap v.1.0.12^80^. Significant differences were calculated by pairwise t-tests with the Benjamini-Hochberg multiple testing procedure.

#### Gene signature scoring

Scoring of gene signatures used the R package `UCell v.2.8.0` to score individual gene signature sets using the Mann-Whitney U statistic^93^. Individual gene sets with p-values for each cell type comparison are available in Table S1. Statistical analysis for each gene signature comparison used estimated marginal means with pairwise comparisons of each cell type between *Ren1*^*c+/-*^;*Ren1*^*cCre*^;*R26R*^*mTmG*^ (*Ren1*^*c*^*WT*) and *Ren1*^*c-/-*^;*Ren1*^*cCre*^;*R26R*^*mTmG*^ (*Ren1*^*c*^*KO*) or *SMMHC*^*CreERT2*^;*R26R*^*tdTomato*^;*Ren1*^*cYFP*^ control versus captopril-treated cells followed by the Benjamini-Hochberg multiple testing procedure.

For the development of the CAAH gene signature, we first merged the individual gene signatures. Then, we iteratively sampled 10 random genes from the superset and scored, using the Mann Whitney U statistic implemented in `UCell`, that gene set for each cell type comparison. We performed 5000 permutations of this procedure and retained the set of 10 genes with the greatest differential score between conditions, either between the different genotypes in the *Ren1*^*c*^ animals, or between captopril-treated or untreated *SMMHC*^*CreERT2*^;*R26R*^*tdTomato*^;*Ren1*^*cYFP*^ animals. Statistical differences were performed as described for each individual gene signature.

### Microscopy

Tissue sections were visualized using a Zeiss Imager M2 microscope equipped with the AxioCam 305 color and AxioCam 506 mono cameras and ApoTome.2 (Zeiss, Oberkochen, Germany). The juxtaglomerular apparatus (JGA) index was calculated as the number of renin-positive JGA/total number of glomeruli and expressed as a percentage^94^. All the image analysis parameters were kept constant between samples.

### 3D visualization of the renal arterial tree

For the tissue-clearing and whole-mount immunostaining of *Ren1*^*c-/-*^;*Ren1*^*cCre*^;*R26R*^*mTmG*^ and *SMMHC*^*CreERT2*^;*R26R*^*tdTomato*^;*Ren1*^*cYFP*^ mice kidneys, we used the updated clear, unobstructed brain/body imaging cocktails and computational analysis (CUBIC) protocol^95^. The mice were anesthetized with tribromoethanol (300 mg/kg) and perfused with 20 mL of PBS and 30 mL of 4% PFA via the left ventricle of the heart. Kidneys were removed, divided into two sections each, and fixed overnight in 4% PFA. All subsequent steps were performed with gentle shaking. The samples were washed with PBS for 6 hours followed by immersion in CUBIC-L (10 wt% N-butyldiethanolamine and 10 wt% Triton X-100) at 45°Cfor 5 days. After washing with PBS for 6 hours, the samples were immersed in CUBIC-R+ [T3741 (45 wt% 2,3-dimethyl-1-phenyl-5-pryrazolone, 30 wt% nicotinamide and 5 wt% N-butyldiethanolamine), Tokyo Chemical Industry, Tokyo, Japan] and diluted 1:1 in water overnight at RT. The samples were then immersed in CUBIC-R+ at RT for 2 days.

Macroscopic whole-mount images were acquired with a light-sheet fluorescence (LSF) microscope (ZEISS Light-sheet7, Zeiss). The samples were imaged in mounting solution (RI:1.520) (M3294, Tokyo Chemical Industry) with Clr Plan-Neofluar 20x/1.0 Corr nd=1.53 detection optics. The voxel resolution was as follows: x = 0.167-0.236 *µ*m, y = 0.167-0.236 *µ*m, z = 0.56-1.0 *µ*m (zoom range*×* 0.36-1.41). The GFP and YFP signals were measured by excitation with 488 nm lasers. The tdTomato signals were measured by excitation with 561 nm lasers.

Three-dimensionally rendered images were visualized and edited with Imaris software (Version 10.0.0, Bitplane, Belfast, United Kingdom). Raw image files obtained from the Zeiss LSF microscope (.czi) were converted into Imaris files (.ims) using Imaris File Converter 10.0.0. Tilescan images were stitched using Imaris Stitcher 10.0.0. The image processing by Imaris software was performed as previously described^57^. The reconstituted 3D images were cropped to a region of interest using the crop function. The snapshot and animation functions were used to capture images and videos, respectively.

### Proteomic analysis of urine samples by mass spectrometry analysis

Aliquots of 100 *µ*L of urine were submitted to the Biomolecular Analysis Facility Core (University of Virginia - Office of Research Core Administration) for proteomic analysis. Protein precipitation was performed by adding-20°Cacetone (1x volume of the sample) and -20°Cmethanol (9*×* volume of the sample) and incubated overnight at -80^*°*^C. Samples were centrifuged 30 min at 4°Cat 10,000*×* g and pellets were washed with 1 mL cold methanol and centrifuged 30 min at 4°Cat 10,000 *×*g. Proteins were solubilized in 50 *µ*L of 50 mM Ammonium Bicarbonate (AmBiC) and the protein concentration of the supernatant was determined using BCA. Twenty micrograms of each sample were transferred into a new, pre-washed tube and reduced using 10 mM Dithiothreitol for 30 min at RT and alkylated with 50 mM Iodacetamide for 30 min at RT. Sample volume was adjusted to a final volume of 100 *µ*L with 50 mM AmBiC and trypsin digestion was performed adding 1 *µ*g of modified trypsin solution overnight at 37^*°*^C. After digestion samples were acidified using 8 *µ*L of formic acid to quench the reaction and submitted to desalt following the BAF Protocol 003 procedure (dx.doi.org/10.17504/protocols.io.36wgqjzmyvk5/v1). The LC-MS system consisted of a Thermo Orbitrap Exploris 480 mass spectrometer system with an Easy Spray ion source connected to a Thermo 3 *µ*m C18 Easy Spray column (through pre-column). Specific parameters for data acquisition (dx.doi.org/10.17504/protocols.io.81wgbxb63lpk/v1) and for database searches (dx.doi.org/10.17504/protocols.io.q26g7p28kgwz/v1) were applied.

### Extraction of hydrophilic metabolites and mass spectrometry analyses

Urine from control and *Ren1*^*c*^*KO* mice (at 1 and 6 months) were submitted to the Biomolecular Analysis Facility Core (BAF Core at the University of Virginia) for metabolome analysis. Hydrophilic metabolites were extracted. After collection, samples were maintained at -80^*°*^C before extraction procedures. Urine samples were defrosted on ice, the osmolarity was measured by the vapor pressure method (Vapro-5600, ELITechGroup, Puteaux, France) and sample volumes were normalized by osmolarity as the *Ren1*^*c*^*KO* animals are incapable of concentrating urine to the same degree as control mice^55^. Samples were lyophilized overnight and resuspended in 100 *µ*L of water. To each tube, 750 *µ*L of -20^*°*^C cold chloroform: methanol (2:1) mixture was added. Tubes were vortexed and shaken vigorously for 30 min at 4^*°*^C in a temperature controlled thermal mixer (Thermo Scientific, Waltham, MA, USA). Four hundred microliters of water were added to induce phase separation and samples were centrifuged at 10,000 rpm for 10 min, at 4^*°*^C. The top aqueous methanolic phase was recovered as metabolite mixture. Metabolite mixtures were stored in Eppendorf tubes, dried under vacuum and stored at -80^*°*^C. Sample preparation for injection, specific parameters for data acquisition of hydrophilic metabolites (dx.doi.org/10.17504/protocols.io.bp2l629ddgqe/v1) and for database searches (dx.doi.org/10.17504/protocols.io.dm6gpzj1jlzp/v1) were applied.

### Statistical Analysis

Data is presented as means *±*SD. Unless stated otherwise, statistical analysis was performed using GraphPad Prism 10 software (GraphPad Software, San Diego, CA) or in R. Comparisons between two groups were performed by two-tailed unpaired Student’s t-test when passing tests for normality and heterogeneity of variance. Comparisons between more than two groups were performed using Welch’s ANOVA when groups had unequal variances. Posthoc comparisons between group means were evaluated using Dunnett’s T3 multiple comparison test or pairwise comparisons of estimated marginal means. P values *<* 0.05 were considered statistically significant.

## Supplemental Figures

**Fig. S1.**
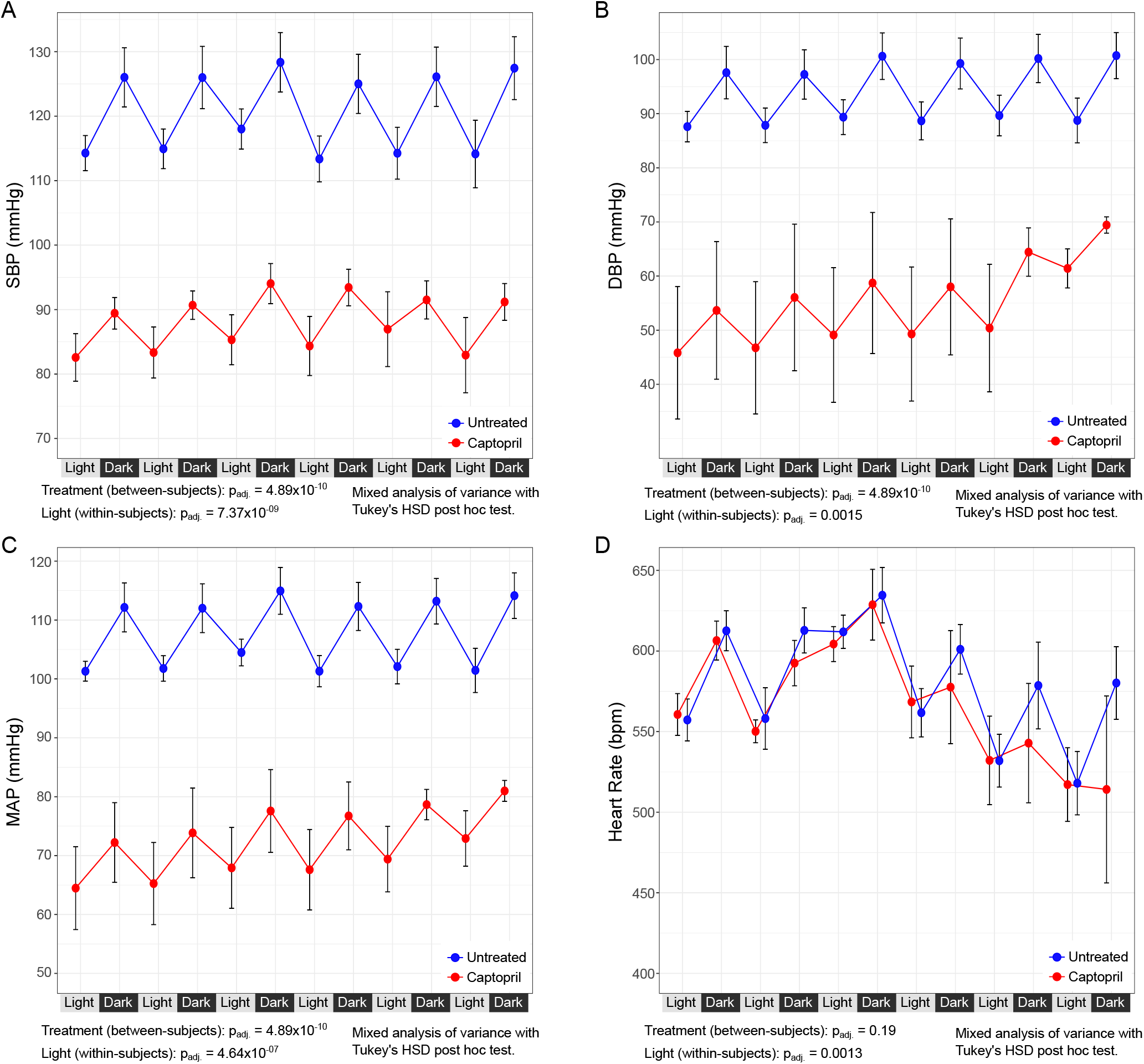
Systolic, diastolic, and mean blood pressure are significantly decreased in captopril-treated SMMHC^CreERT2;^ R26R^tdTomato^;Ren1^cYFP^ mice. Systolic (A), diastolic (B), and mean blood pressure (C) are significantly elevated, irrespective of day or light cycle. (D) Mouse heart rate is not significantly different between captopril treated or untreated animals. Statistical analysis represents the results of a mixed analysis of variance followed by Tukey’s post-hoc test. Video : **Ren1c-lineage cells are localized to the juxtaglomerular area under normal conditions, related to Figure 1**. Three-dimensional visualization of *Ren1*^*c*^-lineage cells (GFP, green) and surrounding background structures (tdTomato, red) in the kidney cortex of a 1-month-old *Ren1*^*c+/-*^;*Ren1*^*cCre*^;*R26R*^*mTmG*^ mouse. Video : **Concentric arterial and arteriolar hypertrophy (CAAH) in *Ren1***^***c***^***KO* mouse, related to Figure 1**. Three-dimensional distribution of *Renin*^*null*^/*Ren1*^*c*^*KO* cells in 1-month-old *Ren1*^*c-/-*^;*Ren1*^*cCre*^;*R26R*^*mTmG*^ mouse. Hypertrophied *Renin*^*null*^ cells accumulate in arterioles, forming CAAH that often encircle the glomeruli. *Renin*^*null*^ cells are also broadly distributed along arcuate arteries. Video : **Three-dimensional morphology of the renal vascular tree and renin cells under normal conditions, related to Figure 1**. Kidney cortex from a 5-month-old *SMMHC*^*CreERT2*^;*R26R*^*tdTomato*^;*Ren1*^*cYFP*^ mouse (control). Vascular smooth muscle is labeled with tdTomato (red), and renin cells (YFP, yellow) are localized exclusively at the tips of afferent arterioles. Video : **Concentric arterial and arteriolar hypertrophy (CAAH) in captopril-treated mouse, related to Figure 1**. Kidney cortex from a 5-month-old *SMMHC*^*CreERT2*^;*R26R*^*tdTomato*^;*Ren1*^*cYFP*^ mouse treated with captopril for 3

**Fig. S2.**
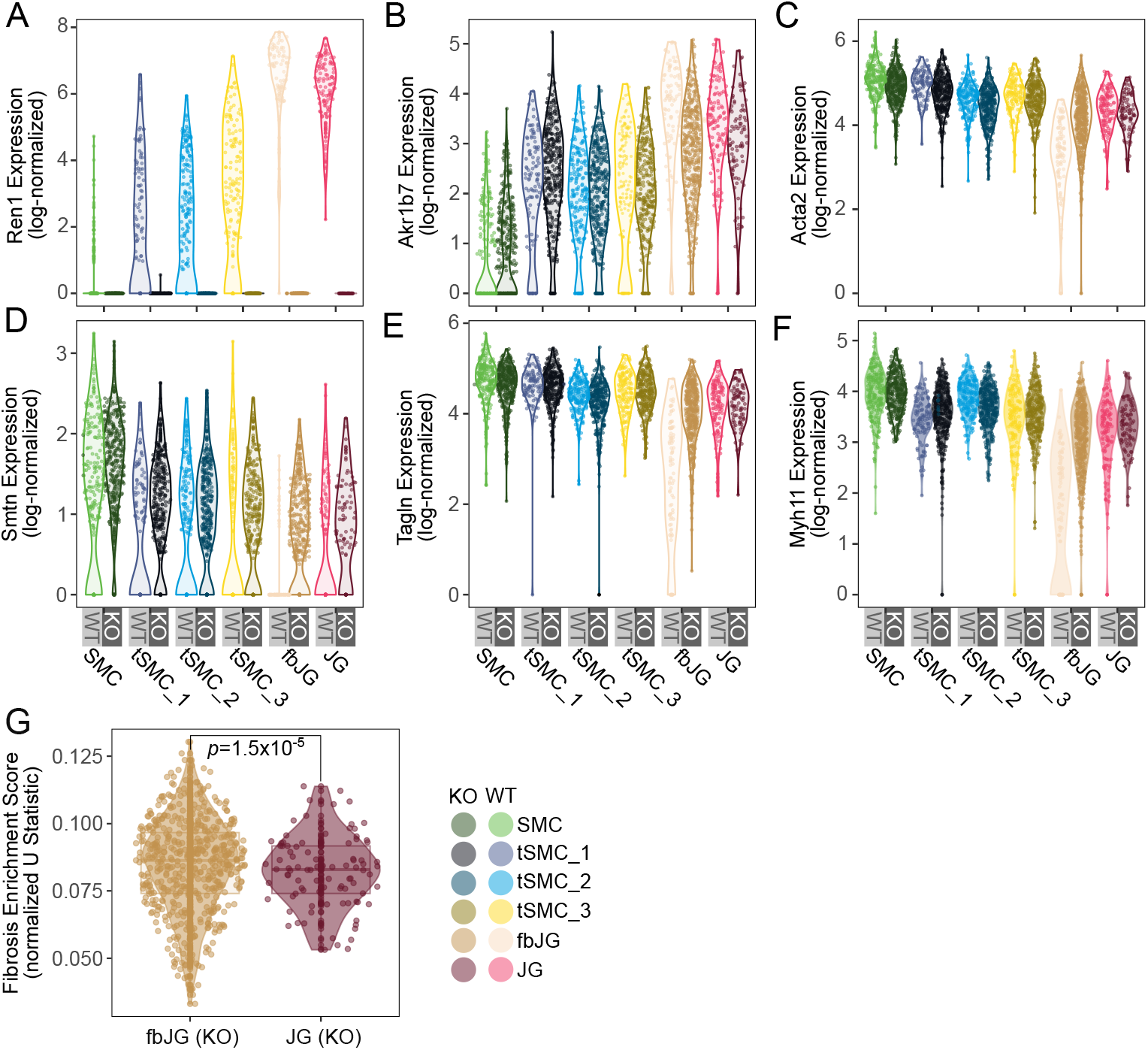
Marker expression for smooth muscle cells (SMC), juxtaglomerular (JG) cells, and fibrotic (fb) cells in renin-lineage cell populations from Ren1^c^KO and Ren1^c^WT mice. (A) Ren1 is most highly expressed in JG populations and is absent, as expected, in Ren1^c^KO populations. (B) Akr1b7, a surrogate JG marker, is expressed in renin-lineage populations with the highest expression in JG cells. SMC marker genes, including Acta2 (C), Smtn (D), Tagln (E), and Myh11 (F) are highly expressed in SMC cells and to a lesser degree in JG cells. (G) A gene signature of fibrosis is elevated in the fibrotic juxtaglomerular cell population (fbJG), leading to the assigned nomenclature. P value represents Welch’s T test.

**Fig. S3.**
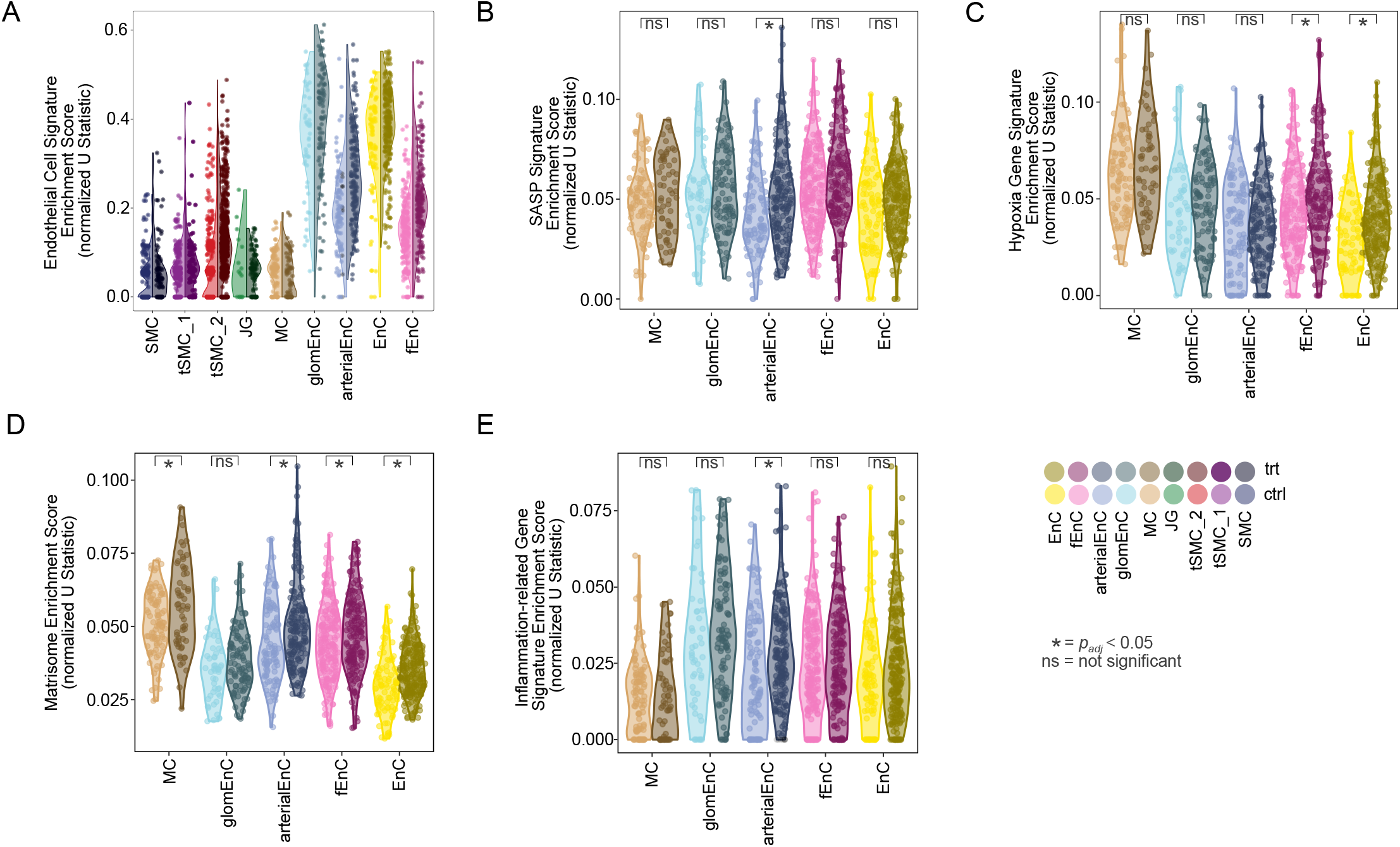
Cell and gene signature analysis in additional SMMHC^CreERT2^;R26R^tdTomato;^ Ren1^cYFP^ mice trajectory populations. The trajectory includes distinct populations of endothelial cells (A). Compared to the primary renin-lineage cell populations, only arterial endothelial cells (arterialEnC) display an enriched signature of the senescence-associated secretory phenotype (B). Both fenestrated (fEnC) and basal endothelial cells (EnC) have an enriched signature of hypoxia in RAS inhibited populations (C). All populations but glomerular endothelial cells (glomEnC) have increased extracellular matrix deposition in RAS inhibited populations (D). Only the arterial endothelial cells have an inflammation-related gene signature in RAS inhibited populations relative to normal (E). (=p*<0.05. Statistical tests represent an estimated marginal means linear model followed by pairwise comparisons between treated (trt) and untreated (ctrl) cell populations with the Benjamini-Hochberg multiple testing procedure correction.) months. The tips of afferent arterioles are thickened in a goblet-like configuration and enveloped by renin cells, forming CAAH. Striated extensions (recruitment) of renin cells are frequently observed along the renal vascular tree (tdTomato, red).

## References

1. (NCD-RisC) NRFC. Worldwide trends in hypertension prevalence and progress in treatment and control from 1990 to 2019: A pooled analysis of 1201 population-representative studies with 104 million participants. Lancet (London, England). 2021;398:957–980.

2. Organization WH. Fact sheet on hypertension. 2023. Available at https://www.who.int/news-room/fact-sheets/detail/hypertension.

3. Falkner B, Gidding SS, Baker-Smith CM, Brady TM, Flynn JT, Malle LM, South AM, Tran AH, Urbina EM, American Heart Association Council on Hypertension; Council on Lifelong Congenital Heart Disease on behalf of the, Young; Council on Kidney in Cardiovascular Disease; Council on Lifestyle HH in the, Health; C, Cardiovascular C on, Nursing S. Pediatric primary hypertension: An underrecognized condition: A scientific statement from the american heart association. Hypertension. 2023;80:e101–e111.

4. Whelton PK, Carey RM, Aronow WS, Casey DEJ, Collins KJ, Dennison Himmelfarb C, DePalma SM, Gidding S, Jamerson KA, Jones DW, MacLaughlin EJ, Muntner P, Ovbiagele B, Smith SCJ, Spencer CC, Stafford RS, Taler SJ, Thomas RJ, Williams KAS, Williamson JD, Wright JTJ. 2017 ACC/AHA/AAPA/ABC/ACPM/AGS/APhA/ASH/ASPC/NMA/PCNA guideline for the prevention, detection, evaluation, and management of high blood pressure in adults: A report of the american college of cardiology/american heart association task force on clinical practice guidelines. Hypertension (Dallas, Tex : 1979). 2018;71:e13–e115.

5. Mancia G, Kreutz R, Brunström M, Burnier M, Grassi G, Januszewicz A, Muiesan ML, Tsioufis K, Agabiti-Rosei E, Algharably EAE, Azizi M, Benetos A, Borghi C, Hitij JB, Cifkova R, Coca A, Cornelissen V, Cruickshank JK, Cunha PG, Danser AHJ, Pinho RM de, Delles C, Dominiczak AF, Dorobantu M, Doumas M, Fernández-Alfonso MS, Halimi J-M, Járai Z, Jelaković B, Jordan J, Kuznetsova T, Laurent S, Lovic D, Lurbe E, Mahfoud F, Manolis A, Miglinas M, Narkiewicz K, Niiranen T, Palatini P, Parati G, Pathak A, Persu A, Polonia J, Redon J, Sarafidis P, Schmieder R, Spronck B, Stabouli S, Stergiou G, Taddei S, Thomopoulos C, Tomaszewski M, Van de Borne P, Wanner C, Weber T, Williams B, Zhang Z-Y, Kjeldsen SE. 2023 ESH guidelines for the management of arterial hypertension the task force for the management of arterial hypertension of the european society of hypertension: Endorsed by the international society of hypertension (ISH) and the european renal association (ERA). Journal of hypertension. 2023;41:1874–2071.

6. Watanabe H, Martini AG, Brown EA, Liang X, Medrano S, Goto S, Narita I, Arend LJ, Sequeira-Lopez Mls, Gomez RA. Inhibition of the renin-angiotensin system causes concentric hypertrophy of renal arterioles in mice and humans. JCI Insight. 2021;6. doi:10.1172/jci.insight.154337.

7. Oka M, Medrano S, Sequeira-Lopez MLS, Gómez RA. Chronic stimulation of renin cells leads to vascular pathology. Hypertension. 2017;70:119–128.

8. Nagai Y, Yamabe F, Sasaki Y, Ishii T, Nakanishi K, Nakajima K, Shibuya K, Mikami T, Akasaka Y, Urita Y, Yamanaka N. A study of morphological changes in renal afferent arterioles induced by angiotensin II type 1 receptor blockers in hypertensive patients. Kidney & blood pressure research. 2020;45:194–208.

9. Almeida LF, Gomez RA, Sequeira-Lopez MLS. Elusive and heterogenous nature of renin cells. Hypertension. 2024;81:203–205.

10. Pentz ES, Moyano MA, Thornhill BA, Lopez MLSS, Gomez RA. Ablation of renin-expressing juxtaglomerular cells results in a distinct kidney phenotype. Am J Physiol Regul Integr Comp Physiol. 2004;286:R474–R483.

11. Lin EE, Pentz ES, Sequeira-Lopez MLS, Gomez RA. Aldo-keto reductase 1b7, a novel marker for renin cells, is regulated by cyclic AMP signal- ing. Am J Physiol Regul Integr Comp Physiol. 2015;309:R576–84.

12. Coll-Bonfill N, Musri MM, Ivo V, Barbera JA, Tura-Ceide O. Transdifferentiation of endothelial cells to smooth muscle cells play an important role in vascular remodelling. American journal of stem cells. 2015;4:13–21.

13. Frid MG, Kale VA, Stenmark KR. Mature vascular endothelium can give rise to smooth muscle cells via endothelial-mesenchymal transdifferentiation: In vitro analysis. Circulation research. 2002;90:1189–96.

14. Sequeira Lopez ML, Pentz ES, Robert B, Abrahamson DR, Gomez RA. Embryonic origin and lin- eage of juxtaglomerular cells. American journal of physiology Renal physiology. 2001;281:F345–56.

15. Sequeira-Lopez MLS, Lin EE, Li M, Hu Y, Sigmund CD, Gomez RA. The earliest metanephric arte- riolar progenitors and their role in kidney vas- cular development. American journal of physiology Regulatory, integrative and comparative physiology. 2015;308:R138–49.

16. Bradshaw AD. The role of SPARC in extracellular matrix assembly. Journal of cell communication and signaling. 2009;3:239–46.

17. Chen S-J, Ning H, Ishida W, Sodin-Semrl S, Takagawa S, Mori Y, Varga J. The early-immediate gene EGR-1 is induced by transforming growth factor-β and mediates stimulation of collagen gene expression*. Journal of Biological Chemistry. 2006;281:21183–21197.

18. Dickinson MG, Kowalski PS, Bartelds B, Borgdorff MAJ, Feen D van der, Sietsma H, Molema G, Kamps JAAM, Berger RMF. A critical role for egr-1 during vascular remodelling in pulmonary arterial hypertension. Cardiovascular research. 2014;103:573–84.

19. Guerquin M-J, Charvet B, Nourissat G, Havis E, Ronsin O, Bonnin M-A, Ruggiu M, Olivera-Martinez I, Robert N, Lu Y, Kadler KE, Baumberger T, Doursounian L, Berenbaum F, Duprez D. Transcription factor EGR1 directs tendon differentiation and promotes tendon repair. The Journal of clinical investigation. 2013;123:3564–76.

20. Hu F, Xue M, Li Y, Jia Y-J, Zheng Z-J, Yang Y-L, Guan M-P, Sun L, Xue Y-M. Early growth response 1 (Egr1) is a transcriptional activator of NOX4 in oxidative stress of diabetic kidney disease. Journal of diabetes research. 2018;2018:3405695.

21. Oguro A, Koyama C, Xu J, Imaoka S. A cellular stress response (CSR) that interacts with NADPH-P450 reductase (NPR) is a new regulator of hypoxic response. Biochemical and Biophysical Research Communications. 2014;445:43–47.

22. Montesi SB, Izquierdo-Garcia D, Dèsogère P, Abston E, Liang LL, Digumarthy S, Seethamraju R, Lanuti M, Caravan P, Catana C. Type i collagen-targeted positron emission tomography imaging in idiopathic pulmonary fibrosis: First-in-human studies. American journal of respiratory and critical care medicine. 2019;200:258–261.

23. Baues M, Klinkhammer BM, Ehling J, Gremse F, Zandvoort MAMJ van, Reutelingsperger CPM, Daniel C, Amann K, Bábíčková J, Kiessling F, Floege J, Lammers T, Boor P. A collagen-binding protein enables molecular imaging of kidney fibrosis in vivo. Kidney international. 2020;97:609–614.

24. Kudrjashova E, Bashtrikov P, Bochkov V, Parfyonova Y, Tkachuk V, Antropova J, Iljinskaya O, Tararak E, Erne P, Ivanov D, Philippova M, Resink TJ. Expression of adhesion molecule t-cadherin is increased during neointima formation in experimental restenosis. Histochemistry and cell biology. 2002;118:281–90.

25. Org E, Eyheramendy S, Juhanson P, Gieger C, Lichtner P, Klopp N, Veldre G, Döring A, Viigimaa M, Sõber S, Tomberg K, Eckstein G, KORA, Kelgo P, Rebane T, Shaw-Hawkins S, Howard P, Onipinla A, Dobson RJ, Newhouse SJ, Brown M, Dominiczak A, Connell J, Samani N, Farrall M, BRIGHT, Caulfield MJ, Munroe PB, Illig T, Wichmann H-E, Meitinger T, Laan M. Genome-wide scan identifies CDH13 as a novel susceptibility locus contributing to blood pressure determination in two european populations. Human Molecular Genetics. 2009;18:2288–2296.

26. Popov VS, Brodsky IB, Balatskaya MN, Balatskiy AV, Ozhimalov ID, Kulebyakina MA, Semina EV, Arbatskiy MS, Isakova VS, Klimovich PS, Sysoeva VY, Kalinina NI, Tkachuk VA, Rubina KA. T-cadherin deficiency is associated with increased blood pressure after physical activity. International journal of molecular sciences. 2023;24.

27. Fujii K, Fujishima Y, Kita S, Kawada K, Fukuoka K, Sakaue T-A, Okita T, Kawada-Horitani E, Nagao H, Fukuda S, Maeda N, Nishizawa H, Shimomura I. Pharmacological HIF-1 activation upregulates extracellular vesicle production synergistically with adiponectin through transcriptional induction and protein stabilization of t-cadherin. Scientific reports. 2024;14:3620.

28. Dong R, Yu J, Yu F, Yang S, Qian Q, Zha Y. IGF-1/IGF-1R blockade ameliorates diabetic kidney disease through normalizing Snail1 expression in a mouse model. American journal of physiology Endocrinology and metabolism. 2019;317:E686–E698.

29. Gurevich E, Segev Y, Landau D. Growth hormone and IGF1 actions in kidney development and function. Cells. 2021;10.

30. Mohebi R, Liu Y, Hansen MK, Yavin Y, Sattar N, Pollock CA, Butler J, Jardine M, Masson S, Heerspink HJL, Januzzi Jr JL. Insulin growth factor axis and cardio-renal risk in diabetic kidney disease: An analysis from the CREDENCE trial. Cardiovascular Diabetology. 2023;22:176.

31. Chowdhury KH, Iqbal MM, Rahman IZ, Saha D, Saha S, Hossain MS, Morshed R, Hassan M, Chaudhury S, Chowdhury M. Nephrology Dialysis Transplantation. 2024;39:gfae069-1393-2902.

32. Kassiri Z, Oudit GY, Kandalam V, Awad A, Wang X, Ziou X, Maeda N, Herzenberg AM, Scholey JW. Loss of TIMP3 enhances interstitial nephritis and fibrosis. Journal of the American Society of Nephrology : JASN. 2009;20:1223–35.

33. Fiorentino L, Cavalera M, Menini S, Marchetti V, Mavilio M, Fabrizi M, Conserva F, Casagrande V, Menghini R, Pontrelli P, Arisi I, D’Onofrio M, Lauro D, Khokha R, Accili D, Pugliese G, Gesualdo L, Lauro R, Federici M. Loss of TIMP3 underlies diabetic nephropathy via FoxO1/STAT1 interplay. EMBO molecular medicine. 2013;5:441–55.

34. Chen H, Chen L, Chen Y, Guo Q, Lin S. Exploring the genetic causal association of TIMP3 on CKD and kidney function: A two-sample mendelian randomization. Frontiers in genetics. 2024;15:1367399.

35. Muendlein A, Brandtner EM, Leiherer A, Geiger K, Heinzle C, Gaenger S, Fraunberger P, Haider D, Saely CH, Drexel H. Evaluation of the association of serum glypican-4 with prevalent and future kidney function. Scientific Reports. 2022;12:10168.

36. Muendlein A, Brandtner EM, Schimpf J, Piribauer M, Geiger K, Heinzle C, Leiherer A, Drexel H, Freistätter O, Neyer U, Zitt E. Evaluation of circulating GPC4 in patients with end-stage kidney disease. Kidney360. 2025.

37. Chen B, Miller AL, Rebelatto M, Brewah Y, Rowe DC, Clarke L, Czapiga M, Rosenthal K, Imamichi T, Chen Y, Chang C-S, Chowdhury PS, Naiman B, Wang Y, Yang D, Humbles AA, Herbst R, Sims GP. S100A9 induced inflammatory responses are mediated by distinct damage associated molecular patterns (DAMP) receptors in vitro and in vivo. PloS one. 2015;10:e0115828.

38. Du L, Chen Y, Shi J, Yu X, Zhou J, Wang X, Xu L, Liu J, Gao J, Gu X, Wang T, Yin Z, Li C, Yan M, Wang J, Yin X, Lu Q. Inhibition of S100A8/A9 ameliorates renal interstitial fibrosis in diabetic nephropathy. Metabolism. 2023;144:155376.

39. Zhang J-N, Gong R, Wang Y-Q, Chong Y, Gu Q-K, Zhao M-B, Huang P, Qi Y-C, Meng X-L, Zhao M-Y. Critical role of S100A9 in sepsis-associated acute kidney injury: Mechanistic insights through pyroptosis pathway modulation. Inflammation. 2024.

40. Cai Y, Wang X-L, Flores AM, Lin T, Guzman RJ. Inhibition of endo-lysosomal function exac-erbates vascular calcification. Scientific Reports. 2018;8:3377.

41. Ruby M, Gifford CC, Pandey R, Raj VS, Sabbisetti VS, Ajay AK. Autophagy as a therapeutic target for chronic kidney disease and the roles of TGF-B1 in autophagy and kidney fibrosis. Cells. 2023;12.

42. Zhang Y-Y, Zhou X-T, Huang G-Z, Liao W-J, Chen X, Ma Y-R. The pro-fibrotic role of autophagy in renal intrinsic cells: Mechanisms and therapeutic potential in chronic kidney disease. Frontiers in cell and developmental biology. 2024;12:1499457.

43. Eleutherio ECA, Silva Magalhães RS, de Araújo Brasil A, Monteiro Neto JR, de Holanda Paranhos L. SOD1, more than just an antioxidant. Archives of Biochemistry and Biophysics. 2021;697:108701.

44. Holthoff JH, Harville Y, Herzog C, Juncos LA, Karakala N, Arthur JM. SOD1 is a novel prognostic biomarker of acute kidney injury follow- ing cardiothoracic surgery. BMC Nephrology. 2023;24:299.

45. LeBleu VS, Teng Y, O’Connell JT, Charytan D, Müller GA, Müller CA, Sugimoto H, Kalluri R. Identification of human epididymis protein-4 as a fibroblast-derived mediator of fibrosis. Nature medicine. 2013;19:227–31.

46. Nakagawa S, Nishihara K, Miyata H, Shinke H, Tomita E, Kajiwara M, Matsubara T, Iehara N, Igarashi Y, Yamada H, Fukatsu A, Yanagita M, Matsubara K, Masuda S. Molecular markers of tubulointerstitial fibrosis and tubular cell damage in patients with chronic kidney disease. PloS one. 2015;10:e0136994.

47. Ihara K, Skupien J, Kobayashi H, Md Dom ZI, Wilson JM, O’Neil K, Badger HS, Bowsman LM, Satake E, Breyer MD, Duffin KL, Krolewski AS. Profibrotic circulating proteins and risk of early progressive renal decline in patients with type 2 diabetes with and without albuminuria. Diabetes care. 2020;43:2760–2767.

48. Zhang L, Liu L, Bai M, Liu M, Wei L, Yang Z, Qian Q, Ning X, Sun S. Hypoxia-induced HE4 in tubular epithelial cells promotes extracellular matrix accumulation and renal fibrosis via NF-κb. The FASEB Journal. 2020;34:2554–2567.

49. Kuo DS, Labelle-Dumais C, Gould DB. COL4A1 and COL4A2 mutations and disease: Insights into pathogenic mechanisms and potential therapeutic targets. Human molecular genetics. 2012;21:R97–110.

50. Sanaei-Ardekani M, Kamal S, Handy W, Alam S, Salaheldin A, Moore A, Movafagh S. Suppression of collagen IV alpha-2 subunit by prolyl hydroxylase domain inhibition via hypoxia-inducible factor-1 in chronic kidney disease. Pharmacology Research & Perspectives. 2021;9:e00872.

51. Jones BA, Gisch DL, Myakala K, Sadiq A, Cheng Y-H, Taranenko E, Panov J, Korolowicz K, Melo Ferreira R, Yang X, Santo BA, Allen KC, Yoshida T, Wang XX, Rosenberg AZ, Jain S, Eadon MT, Levi M. NAD+ prevents chronic kidney disease by activating renal tubular metabolism. JCI insight. 2025;10.

52. Murphy MP, O’Neill LAJ. Krebs cycle reimagined: The emerging roles of succinate and itaconate as signal transducers. Cell. 2018;174:780–784.

53. Zhang W, Lang R. Succinate metabolism: A promising therapeutic target for inflammation, ischemia/reperfusion injury and cancer. Frontiers in cell and developmental biology. 2023;11:1266973.

54. Zhang X, Lyu D, Li S, Xiao H, Qiu Y, Xu K, Chen N, Deng L, Huang H, Wu R. Discovery of a SUCNR1 antagonist for potential treatment of diabetic nephropathy: In silico and in vitro studies. International Journal of Biological Macromolecules. 2024;268:131898.

55. Takahashi N, Lopez MLSS, Cowhig JEJ, Taylor MA, Hatada T, Riggs E, Lee G, Gomez RA, Kim H-S, Smithies O. Ren1c homozygous null mice are hypotensive and polyuric, but heterozygotes are indistinguishable from wild-type. Journal of the American Society of Nephrology : JASN. 2005;16:125–32.

56. Kaverina NV, Kadoya H, Eng DG, Rusiniak ME, Sequeira-Lopez MLS, Gomez RA, Pippin JW, Gross KW, Peti-Peterdi J, Shankland SJ. Tracking the stochastic fate of cells of the renin lineage after podocyte depletion using multicolor reporters and intravital imaging. PloS one. 2017;12:e0173891.

57. Medrano S, Yamaguchi M, Almeida LF de, Smith JP, Yamaguchi H, Sigmund CD, Sequeira-Lopez MLS, Gomez RA. An efficient inducible model for the control of gene expression in renin cells. American journal of physiology Renal physiology. 2024;327:F489–F503.

58. Pentz ES, Lopez MLSS, Cordaillat M, Gomez RA. Identity of the renin cell is mediated by cAMP and chromatin remodeling: An in vitro model for studying cell recruitment and plasticity. Am J Physiol Heart Circ Physiol. 2008;294:H699–707.

59. Scarfe L, Schock-Kusch D, Ressel L, Friedemann J, Shulhevich Y, Murray P, Wilm B, Caestecker M de. Transdermal measurement of glomerular filtration rate in mice. Journal of visualized experiments : JoVE. 2018. doi:10.3791/58520.

60. Gomez RA, Pentz ES, Jin X, Cordaillat M, Sequeira Lopez MLS. CBP and p300 are essential for renin cell identity and morphological integrity of the kidney. American journal of physiology Heart and circulatory physiology. 2009;296:H1255–62.

61. Magaletta ME, Lobo M, Kernfeld EM, Aliee H, Huey JD, Parsons TJ, Theis FJ, Maehr R. Integra- tion of single-cell transcriptomes and chromatin landscapes reveals regulatory programs driving pharyngeal organ development. Nat Commun. 2022;13:457.

62. Dobin A, Davis CA, Schlesinger F, Drenkow J, Zaleski C, Jha S, Batut P, Chaisson M, Gingeras TR. STAR: Ultrafast universal RNA-seq aligner. Bioinformatics. 2013;29:15–21.

63. Stuart T, Butler A, Hoffman P, Hafemeister C, Papalexi E, Mauck WM 3rd, Hao Y, Stoeckius M, Smibert P, Satija R. Comprehensive integration of single-cell data. Cell. 2019;177:1888–1902.e21.

64. Hao Y, Stuart T, Kowalski MH, Choudhary S, Hoffman P, Hartman A, Srivastava A, Molla G, Madad S, Fernandez-Granda C, Satija R. Dictio- nary learning for integrative, multimodal and scalable single-cell analysis. Nat Biotechnol. 2024;42:293–304.

65. McCarthy DJ, Campbell KR, Lun ATL, Wills QF. Scater: Pre-processing, quality control, normalization and visualization of single-cell RNA-seq data in r. Bioinformatics (Oxford, England). 2017;33:1179–1186.

66. Ziegenhain C, Vieth B, Parekh S, Reinius B, Guillaumet-Adkins A, Smets M, Leonhardt H, Heyn H, Hellmann I, Enard W. Comparative anal- ysis of single-cell RNA sequencing methods. Mol Cell. 2017;65:631–643.e4.

67. Patil A, Patil A. CellKb immune: A manually curated database of mammalian hematopoietic marker gene sets for rapid cell type identification. bioRxiv. 2022. doi:10.1101/2020.12.01.389890.

68. Cao J, Cusanovich DA, Ramani V, Aghamirzaie D, Pliner HA, Hill AJ, Daza RM, McFaline-Figueroa JL, Packer JS, Christiansen L, Steemers FJ, Adey AC, Trapnell C, Shendure J. Joint pro- filing of chromatin accessibility and gene expression in thousands of single cells. Science. 2018;361:1380–1385.

69. Park J, Shrestha R, Qiu C, Kondo A, Huang S, Werth M, Li M, Barasch J, Suszták K. Single-cell transcriptomics of the mouse kidney reveals potential cellular targets of kidney disease. Science (New York, NY). 2018;360:758–763.

70. Combes AN, Phipson B, Lawlor KT, Dorison A, Patrick R, Zappia L, Harvey RP, Oshlack A, Little MH. Single cell analysis of the developing mouse kidney provides deeper insight into marker gene expression and ligand-receptor crosstalk. Development (Cambridge, England). 2019;146. doi:10.1242/dev.178673.

71. Stuart T, Srivastava A, Madad S, Lareau CA, Satija R. Single-cell chromatin state analysis with signac. Nature methods. 2021;18:1333–1341.

72. Hafemeister C, Satija R. Normalization and variance stabilization of single-cell RNA-seq data using regularized negative binomial regression. Genome Biol. 2019;20:296.

73. Haghverdi L, Lun ATL, Morgan MD, Marioni JC. Batch effects in single-cell RNA-sequencing data are corrected by matching mutual nearest neighbors. Nature Biotechnology. 2018;36:421–427.

74. Korsunsky I, Nathan A, Millard N, Raychaudhuri nS. Presto: Fast functions for differential expression using wilcox and AUC. ; 2025. Available at https://github.com/immunogenomics/presto.

75. Schep AN, Wu B, Buenrostro JD, Greenleaf WJ. chromVAR: Inferring transcription-factor-associated accessibility from single-cell epigenomic data. Nat Methods. 2017;14:975.

76. Trapnell C, Cacchiarelli D, Grimsby J, Pokharel P, Li S, Morse M, Lennon NJ, Livak KJ, Mikkelsen TS, Rinn JL. The dynamics and regulators of cell fate decisions are revealed by pseudotemporal ordering of single cells. Nature biotechnology. 2014;32:381–386.

77. Qiu X, Mao Q, Tang Y, Wang L, Chawla R, Pliner HA, Trapnell C. Reversed graph embedding resolves complex single-cell trajectories. Nature Methods. 2017;14:979–982.

78. Cao J, Spielmann M, Qiu X, Huang X, Ibrahim DM, Hill AJ, Zhang F, Mundlos S, Christiansen L, Steemers FJ, Trapnell C, Shendure J. The single-cell transcriptional landscape of mammalian organogenesis. Nature. 2019;566:496–502.

79. Finak G, McDavid A, Yajima M, Deng J, Gersuk V, Shalek AK, Slichter CK, Miller HW, McElrath MJ, Prlic M, Linsley PS, Gottardo R. MAST: A flexible statistical framework for assessing transcriptional changes and characterizing heterogeneity in single-cell RNA sequencing data. Genome Biology. 2015;16:278.

80. Kolde R. Pheatmap: Pretty heatmaps. ; 2019. Available at https://CRAN.R-project.org/package=pheatmap.

81. Jiang J. IReNA: IReNA. ; 2024. Available at https://github.com/jiang-junyao/IReNA.

82. Kassambara A, Mundt F. Factoextra: Extract and visualize the results of multivariate data analyses. ; 2020. Available at http://www.sthda.com/english/rpkgs/factoextra.

83. Huynh-Thu VA, Irrthum A, Wehenkel L, Geurts P. Inferring regulatory networks from expression data using tree-based methods. PloS one. 2010;5.

84. Aibar S, González-Blas CB, Moerman T, Huynh-Thu VA, Imrichova H, Hulselmans G, Rambow F, Marine J-C, Geurts P, Aerts J, Oord J van den, Atak ZK, Wouters J, Aerts S. SCENIC: Single-cell regulatory network inference and clustering. Nature Methods. 2017;14:1083–1086.

85. Schep A. Motifmatchr: Fast motif matching in r. ; 2024. doi:10.18129/B9.bioc.motifmatchr.

86. Shannon P, Markiel A, Ozier O, Baliga NS, Wang JT, Ramage D, Amin N, Schwikowski B, Ideker T. Cytoscape: A software environment for integrated models of biomolecular interaction networks. Genome research. 2003;13:2498–504.

87. Fleck JS, Jansen SMJ, Wollny D, Zenk F, Seimiya M, Jain A, Okamoto R, Santel M, He Z, Camp JG, Treutlein B. Inferring and perturbing cell fate regulomes in human brain organoids. Nature. 2023;621:365–372.

88. McLean CY, Bristor D, Hiller M, Clarke SL, Schaar BT, Lowe CB, Wenger AM, Bejerano G. GREAT improves functional interpretation of cis-regulatory regions. Nat Biotechnol. 2010;28:495–501.

89. Tanigawa Y, Dyer ES, Bejerano G. WhichTF is functionally important in your open chromatin data? PLoS Comput Biol. 2022;18:e1010378.

90. Subramanian A, Tamayo P, Mootha VK, Mukherjee S, Ebert BL, Gillette MA, Paulovich A, Pomeroy SL, Golub TR, Lander ES, Mesirov JP. Gene set enrichment analysis: A knowledge-based approach for interpreting genome-wide expression profiles. Proceedings of the National Academy of Sciences. 2005;102:15545–15550.

91. Castanza AS, Recla JM, Eby D, Thorvaldsdóttir H, Bult CJ, Mesirov JP. Extending support for mouse data in the molecular signatures database (MSigDB). Nature Methods. 2023;20:1619–1620.

92. Davis D, Wizel A, Drier Y. Accurate estimation of pathway activity in single cells for clustering and differential analysis. Genome research. 2024;34:925–936.

93. Andreatta M, Carmona SJ. UCell: Robust and scalable single-cell gene signature scoring. Computational and structural biotechnology journal. 2021;19:3796–3798.

94. Guessoum O, Zainab M, Sequeira-Lopez MLS, Gomez RA. Proliferation does not contribute to murine models of renin cell recruitment. Acta Physiol. 2020;230. doi:10.1111/apha.13532.

95. Tainaka K, Murakami TC, Susaki EA, Shimizu C, Saito R, Takahashi K, Hayashi-Takagi A, Sekiya H, Arima Y, Nojima S, Ikemura M, Ushiku T, Shimizu Y, Murakami M, Tanaka KF, Iino M, Kasai H, Sasaoka T, Kobayashi K, Miyazono K, Morii E, Isa T, Fukayama M, Kakita A, Ueda HR. Chemical landscape for tissue clearing based on hydrophilic reagents. Cell reports. 2018;24:2196–2210.e9.

